# Building a mechanistic mathematical model of hepatitis C virus entry

**DOI:** 10.1101/336495

**Authors:** Mphatso Kalemera, Dilyana Mincheva, Joe Grove, Christopher J. R. Illingworth

## Abstract

The mechanism by which hepatitis C virus (HCV) gains entry into cells is a complex one, involving a broad range of host proteins. Entry is a critical phase of the viral lifecycle, and a potential target for therapeutic or vaccine-mediated intervention. However, the mechanics of HCV entry remain poorly understood. Here we describe a novel computational model of viral entry, encompassing the relationship between HCV and the key host receptors CD81 and SR-B1. We conduct experiments to thoroughly quantify the influence of an increase or decrease in receptor availability upon the extent of viral entry. We use these data to build and parameterise a mathematical model, which we then validate by further experiments. Our results are consistent with sequential HCV-receptor interactions, whereby initial interaction between the HCV E2 glycoprotein and SR-B1 facilitates the accumulation CD81 receptors, leading to viral entry. However, we also demonstrate that a small minority of virus can achieve entry in the absence of SR-B1. Our model estimates the impact of the different obstacles that viruses must surmount to achieve entry; among virus particles attaching to the cell surface, 20-35% accumulate sufficient CD81 receptors, of these 4-8% then complete the subsequent steps to achieve productive infection. Furthermore, we make estimates of receptor stoichiometry; between 3 and 6 CD81 receptors are likely to be required to achieve viral entry. Our model provides a tool to investigate the entry characteristics of HCV variants and outlines a framework for future quantitative studies of the multi-receptor dynamics of HCV entry.

## Introduction

HCV can establish a lifelong infection and is a leading cause of liver failure, resulting in 350,000-700,000 deaths annually [1]. A molecular understanding of the intermediate stages of viral replication (e.g. RNA replication, protein processing) has allowed the development of potent antivirals capable of curing *>* 95% of HCV infections [2]. However, of the *∼* 70 million HCV positive individuals, the majority do not know they are infected; consequently transmission rates remain high and may even be increasing [3]. A prophylactic vaccine is the greatest unmet need in our response to HCV; progress towards this goal would be expedited by a molecular understanding of virus entry.

HCV has an enveloped particle that enters via clathrin-mediated endocytosis and low-pH triggered fusion, this process is driven by the glycoproteins E1 and E2 [4]. The molecular events leading up to fusion are only partially understood but are thought to involve at least five essential host factors: CD81, scavenger receptor B1 (SR-B1), epidermal growth factor receptor (EGFR), claudin-1 and occludin [5-9]. However, clear evidence of direct virus-receptor interaction has, at present, only been found for two of these factors: CD81 and SR-B1. The N-terminal hyper-variable region-1 (HVR-1) of the major glycoprotein, E2, forms a linear binding site that interacts with SR-B1 [6,10]. Whereas the CD81 binding site is composed of discontinuous protein domains that are bought together in the tertiary structure of E2 and interact with the large extracellular loop of CD81 [5,11]. Strain specific and broadly neutralising antibodies, which develop during natural infection, act by blocking E2-SR-B1/CD81 interactions [12-15]. A deeper understanding of E2 function and its interactions with SR-B1 and CD81 is likely to aid the design of candidate B-cell targeted vaccines.

In common with other viruses [16], HCV entry proceeds in a stepwise fashion, combining three phases. These interactions initiate a co-ordinated series of molecular events that ultimately unlock the penetration potential of the virus particle, allowing viral genetic material to be delivered across a cellular membrane. Viral entry thus proceeds in a stepwise fashion, combining three phases. During attachment, viral particles encounter cells via passive diffusion, and attach via multiple low-affinity, low-specificity interactions. During engagement, attached viral particles undergo two-dimensional diffusion across the cell surface, resulting in chance encounters with and binding to bone-fide viral receptors. Finally, viral penetration occurs, leading to the delivery of viral genetic material across the cellular membrane. The whole process is time-limited by the intrinsic instability of the viral particle; entry must be achieved before the virus has passed its ‘expiration date’.

The defined stepwise nature of virus entry makes this process highly amenable to mathematical modelling. Indeed, mathematical modelling has a long history of generating insight into processes of viral infection [17]. For example, a model of receptor expression and viral entry was combined with experimental data to show that binding to more than one Tva800 molecule is required for Avian sarcoma leukosis virus to achieve entry [18].

Concerning HCV, modelling approaches have been used to investigate the basic evolutionary properties of the virus [19], its spread between cells [20], and the effectiveness of a variety of viral therapies [21-23]. Previous modelling studies of HCV entry have utilised a model whereby contact between a virus particle and the cell membrane leads to the formation of a stochastic number of E2-CD81 complexes, with viral entry occurring on the formation of a given number of complexes. This model has been used to estimate that between 1 and 13 complexes are required for entry [24], and further developed to explain changes in the cellular expression of claudin-1 following HCV infection [25].

Here we combine experimental data with a novel mathematical model to evaluate the relationship between the HCV protein E2 and the cellular receptors SR-B1 and CD81. Using basic virological assays we measured attachment and receptor engagement by HCV. These observations, combined with previous literature, allowed us to construct a putative minimal model for HCV entry, which we parameterised using the collected experimental data. Our model supports the notion of sequential receptor interactions by HCV, with initial engagement of SR-B1 likely performing a priming function to enhance CD81 interactions. Our model is robust, achieving an excellent fit with our ‘training set’ of experimental data and successfully predicting the outcome of further experiments. The model offers estimates of receptor stoichiometry and the efficiency of HCV entry, and provides a tool for future investigations of E2 glycoprotein function.

## Results

To assess HCV viral entry we use a combined experimental and mathematical approach, first of all conducting experiments to assess the role of CD81 and SR-B1 in viral entry before using this data to build and refine a mathematical model. To achieve this we exploited the HCVcc system; HCV particles generated in vitro were used to infect human hepatoma cell lines (Huh-7.5 or Huh-7). This system is tractable and manipulable, and generates highly reproducible data; it is also a good model of authentic HCV found in infected patients [26,27].

### Measurement of viral attachment

A virus attachment assay showed that only a minority of virus particles used in our experimental setup attached to Huh-7.5 cells. Viral inoculum was added to wells of an assay plate containing human hepatoma cells (Huh-7.5 or Huh-7). After five hours the number of virus particles associated with the cells was evaluated by qPCR quantification of genome copy numbers (Figure 1). Wells containing human hepatoma cells adsorbed significantly more virus than empty control wells (*∼* 17,000 RNA copies, compared to *∼* 6000); we interpret the difference between these values as representing true levels of virus attachment (i.e. *∼* 11,000 particles). To investigate the potential role of entry receptors in attachment, we also quantified the association of particles with Huh-7 cells in which SR-B1 or CD81 had been genetically ablated by CRISPR Cas9 gene editing. We observed no defect in virus attachment to these cells when compared to parental Huh-7 cells; this is in agreement with a previous study and is consistent with the notion of virus attachment being largely independent of receptor engagement [28-30]. From our measurements we deduced that only *∼* 5% of the experimental inoculum attached to the cells. This apparent bottleneck is likely due to the limited speed of virus particles diffusing in the inoculum volume (100 *μl*); in our setup the majority of virus particles in a well are unlikely to even encounter a cell [31].

**Figure 1.**
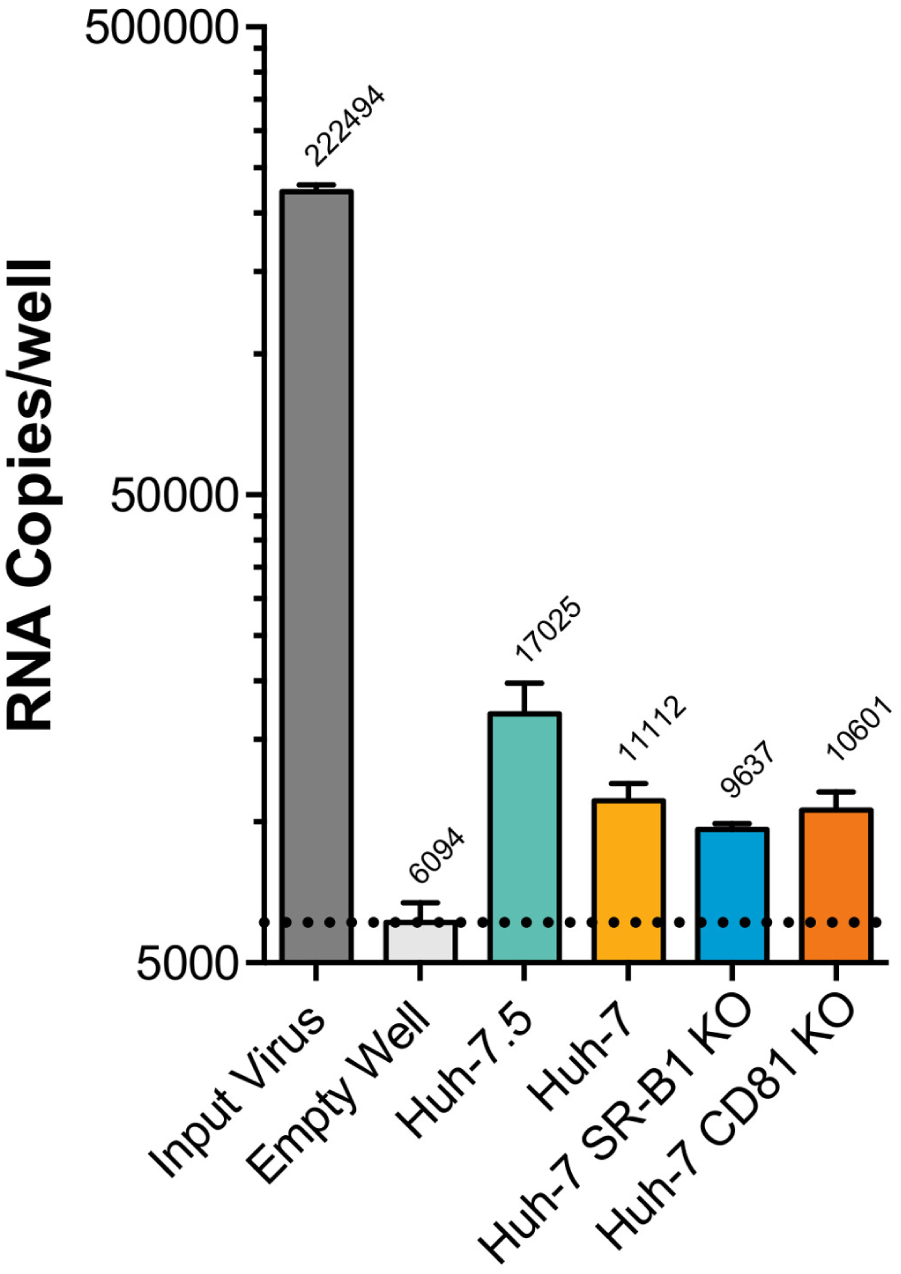
A minority of input virus particles attach to target cells. HCV was inoculated in to replicate wells of a 96 well plate containing the specified cell lines. After five hours the wells were washed with PBS and bound HCV was quantified by qPCR. Each well received *>* 200,000 RNA copies, in an empty well *<* 3% of these remain after washing. Wells containing cells bound significantly more virus than empty wells (p=0.008), however, there was no significant difference between the different cell types (unpaired t-test, GraphPad Prism). The data suggests that *∼* 5% (*∼* 11,000 copies) of the input HCV inoculum binds to Huh-7.5 cells. Mean values of *n >* 3 independent experiments are shown. Error bars indicate the standard error of the mean. The dashed line indicates the background level of virus binding in empty control wells.

### Receptor availability and viral entry

Antibody-mediated blockade experiments showed a decrease in viral entry with decreasing availability of the CD81 and SR-B1 receptors. The basic workflow of our infection assay is summarised in Supporting Figure S1. Huh-7.5 cells were pre-incubated with anti-SR-B1 or anti-CD81 antibodies that are capable of inhibiting E2-receptor interaction and preventing infection [31-33]. The blockaded cells were then challenged with HCV and infectivity was assessed after 48 hours. In a parallel assay plate, antibody binding was assessed by fluorescence microscopy, allowing us to evaluate receptor availability during the virus challenge. A serial dilution of each antibody revealed saturation of binding at high concentrations, suggesting complete receptor blockade; i.e. no receptors were available for entry (Figure 2A,B). Our results show a clear difference between the two receptors; while an absolute blockade of CD81 completely prevented infection, a similar blockade of SR-B1 allowed a small proportion of viruses to achieve infection (Figure 2C). These results suggest that CD81 is absolutely necessary for HCV entry, consistent with many reports [8,26,34,35], while SR-B1 is not strictly necessary for entry to occur. Further support for this conclusion was obtained from measurements of HCV infection of receptor knock-out cells (Supporting Figure S2); modest infection occurred in SR-B1 KO cells, whereas no infection was detected in the absence of CD81. This indicates that a small proportion of virus particles achieve entry in an SR-B1 independent manner [8,26,35].

**Figure 2.**
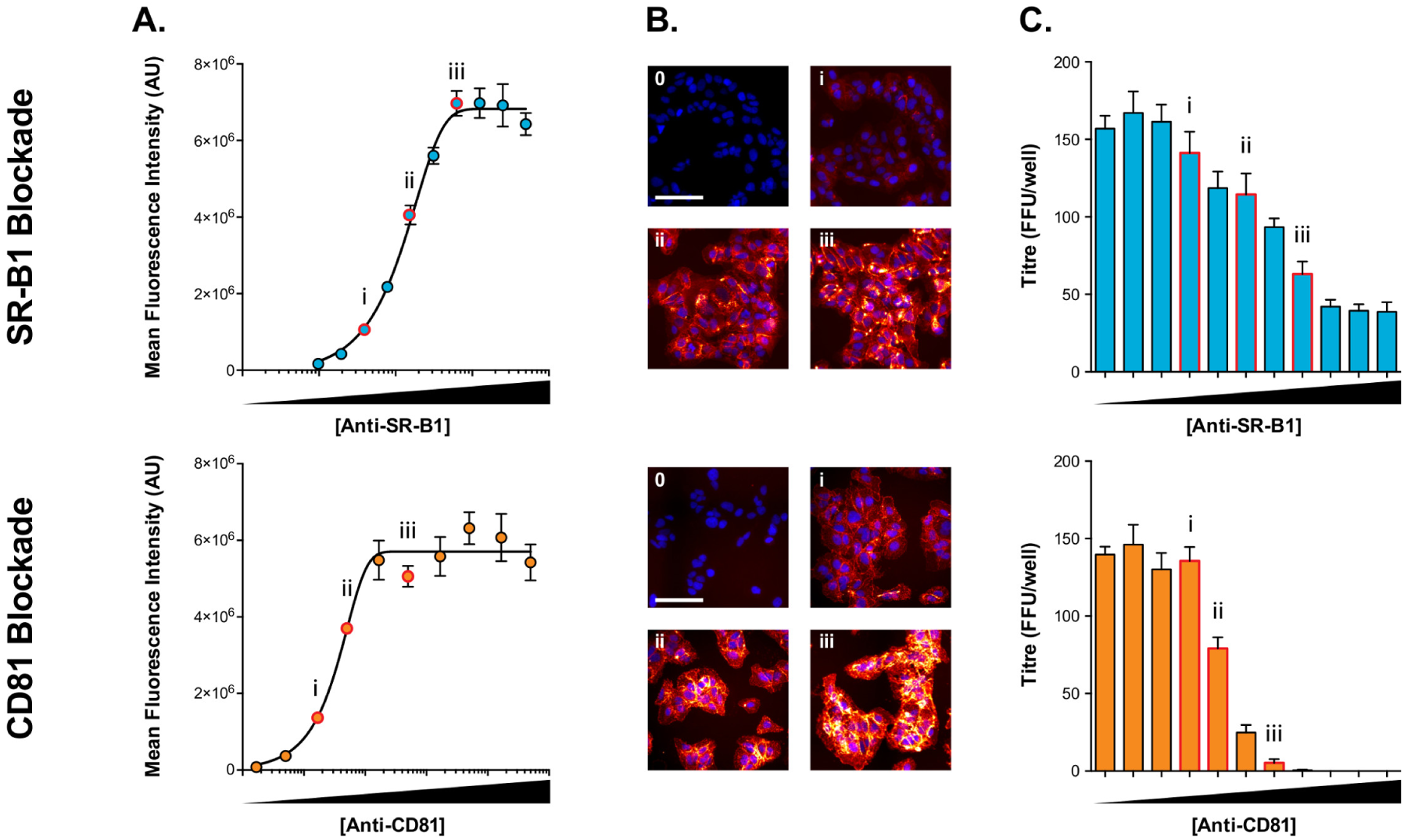
Receptor blockade reveals an SR-B1 independent mode of infection. Huh-7.5 cells were pre-incubated with a dilution series of rabbit anti-SR-B1 serum or anti-CD81 monoclonal supernatant and then challenged with HCVcc. **A.** Binding of anti-receptor antibodies was quantified using fluorescence microscopy, receptor saturation was achieved at high antibody concentrations. Data points represent mean values of *n* = 3 replicates, error bars indicate standard error of the mean. Data was fitted using a sigmoidal curve in GraphPad Prism. **B.** Example fluorescent micrographs of labelled cells, the images are representative of the annotated data points in A. Scale bar 50*μ*m. **C.** HCV infectious titre, expressed as foci per well, in receptor blockaded Huh-7.5 cells, annotated as in A. andB. Complete blockade of CD81 prevents HCV infection, whereas a minority of infection prevails in SR-B1 blockaded cells. Bars represent mean values of *n* = 4 technical replicates, error bars indicate the standard error of the mean. Data from one representative experiment is shown, subsequent modelling was performed with *n* = 3 independent experiments.

Over-expression of SR-B1 and CD81 in cells showed an increase in viral entry with increasing receptor availability. Lentiviral vectors were used to introduce additional receptor coding genes to Huh-7.5 cells. These cells were then challenged with virus as described above (Supporting Figure S1). Parallel plates were fixed and stained for fluorescence microscopy, allowing us to evaluate receptor availability during the virus challenge. Lentiviral transduction resulted in up to four times greater expression than parental cells (Figure 3A). In addition to the receptor coding sequence, the lentiviral vectors also encode GFP, expressed from an additional promoter. Therefore, as an independent measure of transduction, we evaluated GFP expression (Supporting Figure S3); this revealed that all cells were expressing comparable levels of GFP, suggesting homogenous transduction. This is important for unambiguous interpretation of the infectivity data. Whilst infection levels were not directly proportionate to receptor expression, lentiviral transduction resulted in a significant, and dose responsive, increase in HCV infection (Figure 3C). Taken together, the blockade and over-expression data indicate that SR-B1/CD81 availability limits HCV infection.

**Figure 3.**
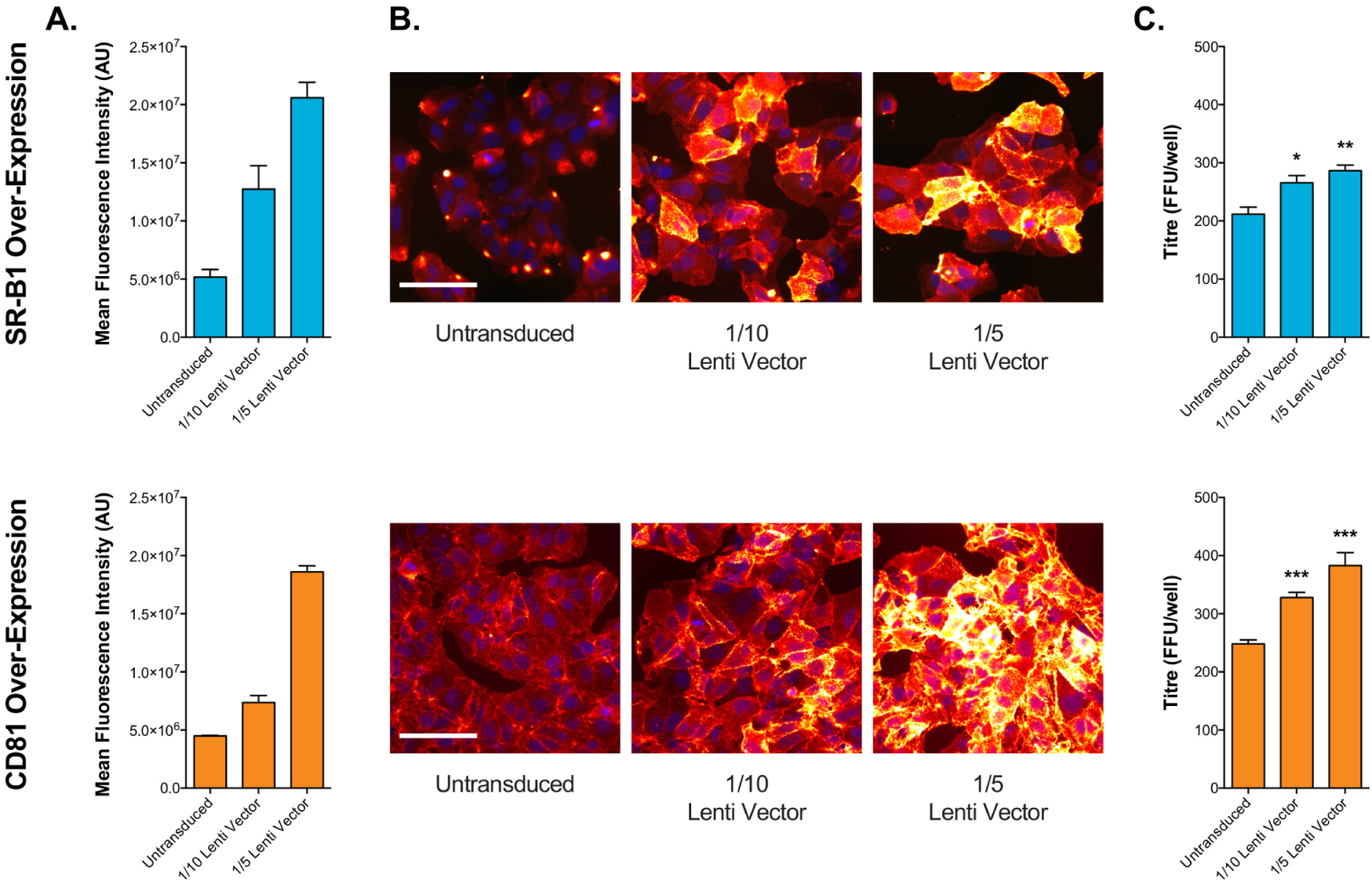
Receptor over-expression enhances HCV infection. Huh-7.5 cells were transduced with lentiviral vectors expressing exogenous human SR-B1 or CD81 and then challenged with HCVcc. **A.** Receptor over-expression was quantified using fluorescence microscopy. Bars represent mean values of *n* = 4 technical replicates, error bars indicate standard error of the mean. **B.** Example fluorescent micrographs of cells over-expressing receptor. Scale bar 50*μ*m. **C.** HCV infectious titre, expressed as foci per well, in receptor over-expressing Huh-7.5 cells. Bars represent mean values of *n* = 4 technical replicates, error bars indicate standard error of the mean. Data from one representative experiment is shown, subsequent modelling was performed with *n* = 3 independent experiments. Significance testing performed against untransduced controls in GraphPad Prism, unpaired t-test, * p*≤*0.01 **p*≤* 0.001 ***p*≤* 0.0001.

### Assessment of virus-receptor interactions

Soluble E2 binding assays suggested that the interaction affinity between E2 and SR-B1 is greater than that between E2 and CD81. Experiments were conducted to measure the binding of recombinant soluble E2 glycoprotein (sE2) to cell surface expressed receptor. This provides a direct measure of virus-receptor interaction, albeit in an artificial system where the components are not presented in their native context.

Chinese hamster ovary (CHO) cells do not bind HCV E2 glycoprotein, however, sE2 binding can be conferred by introduction of exogenous human SR-B1 or CD81; this can be measured by flow cytometry. We treated CHO cells with the receptor + GFP lentiviral vectors (as described above). Antibody staining revealed high levels of receptor expression in GFP positive cells (Supporting Figure S4), with no receptor expression in the minority population of untransduced GFP negative cells. Saturation of binding was achieved at high concentrations of anti-receptor antibody; the intensity of fluorescent signal suggested comparable expression levels for SR-B1 and CD81 (Figure 4A, B). In parallel samples, cells were also incubated with a serial dilution of sE2; example raw fluorescent intensity data is provided in (Supporting Figure S5). sE2 binding to CHO-SR-B1 cells was robust (Figure 4C, D), whereas binding to CHO-CD81 cells was poor. Given the weak binding of sE2 to CHO-CD81 cells, further demonstration of the specificity of sE2 binding to these cells is shown in Supporting Figure S6.

**Figure 4.**
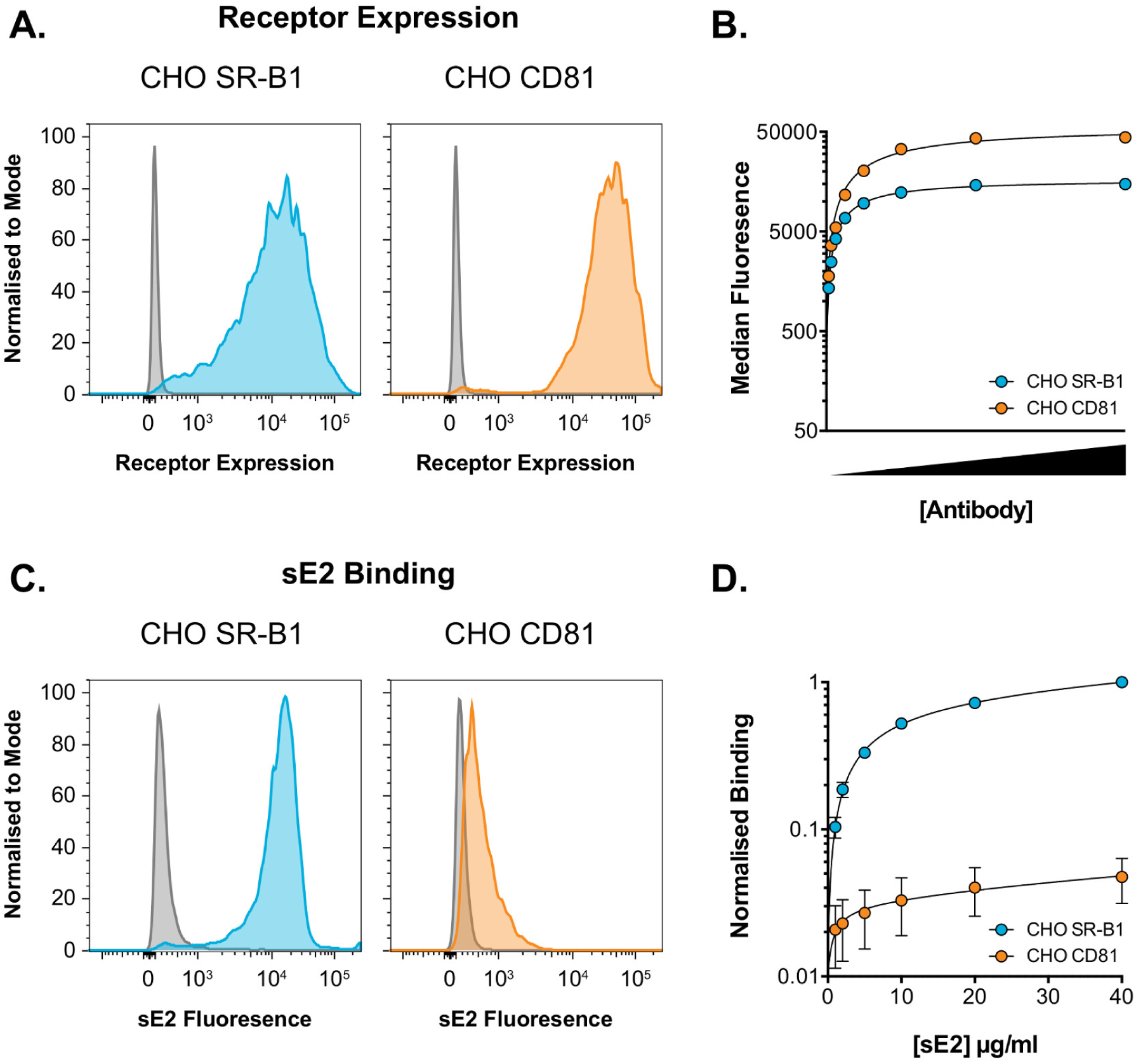
Soluble E2 binds more readily to SR-B1 than CD81. CHO cells were transduced with lentiviral vectors expressing human SR-B1 or CD81 and then used in a sE2-binding assay. **A.** Representative flow cytometry histograms of cell surface expression of HCV receptor in CHO cells, grey plots are untransduced control cells. **B.** Transduced CHO cells were incubated with a serial dilution of anti-receptor antibody. Antibody binding was quantified by flow cytometry; high concentrations of antibody were able to saturate receptor. Data from one representative experiment is shown, data points represent mean values of *n* = 2 technical replicates, error bars indicate standard error of the mean. **C.** Representative flow cytometry histograms of sE2 binding to CHO cells expressing HCV receptor, grey plots are untransduced control cells. **D.** Transduced CHO cells were incubated with a serial dilution of sE2. sE2 binding was quantified by flow cytometry, high concentrations of sE2 approached saturation of binding. Data is normalised to 40*μ* g/ml sE2 binding to CHO SR-B1 cells. Data points represent mean values of *n* = 3 independent experiments, error bars indicate standard error of the mean. Data was fitted using a one-site binding curve in GraphPad Prism.

CD81 and SR-B1 receptors had similar expression in CHO cells (Figure 4B). As such, the binding curves obtained from this experiment (Supporting Figure S5 and Figure 4D) suggest that the intrinsic level of sE2 binding to SR-B1 is up to 20 times greater than that to CD81. However, these observations must be interpreted with care: this assay uses a soluble truncated form of E2, which is devoid of its partner glycoprotein E1 and is no longer presented on the surface of a virus particle. As such, it is probably inappropriate to extract specific values from this data on the relative binding capacity of E2 to either receptor. Taking a conservative interpretation of our data, we concluded simply that HCV is better able to bind to SR-B1 than to CD81, exploring this relationship further during modelling.

### Mathematical model formulation

A mathematical model was used to explain the effect of CD81 and SR-B1 availability upon viral entry. Drawing on the experimental data presented here and past literature [7,36-39] we constructed a minimally complex mechanistic model of HCV receptor engagement (Figure 5A). In our model, once attached to the cell surface, a virus particle must acquire a sufficient number of CD81 receptors to achieve entry; without sufficient CD81, entry cannot occur. The number of CD81 receptors required for entry is unknown, but is characterised by a parameter, *r*, in our model. Models with different number of required receptors were compared, evaluating the number of receptors likely to be needed for viral entry.

**Figure 5.**
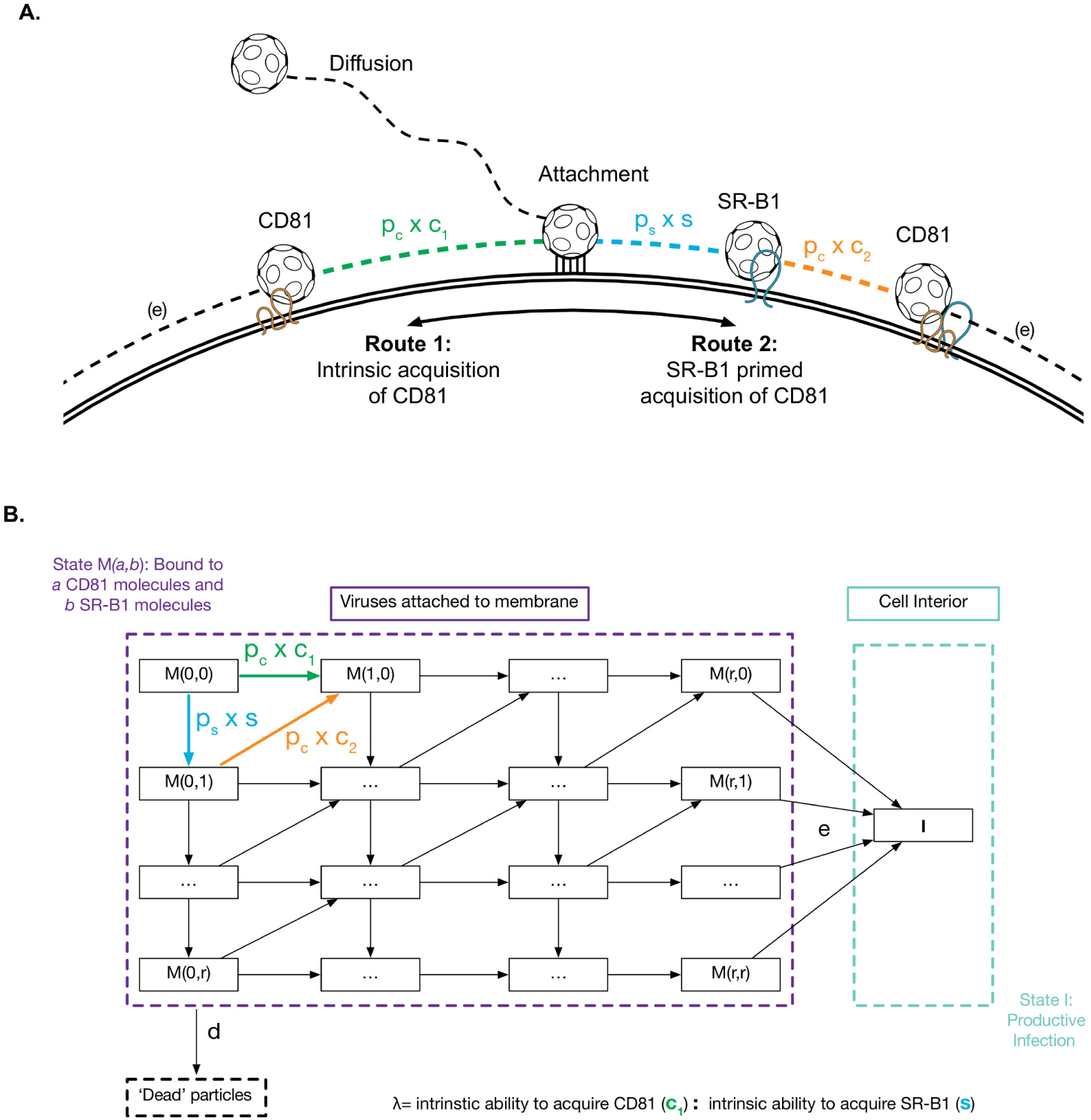
Building a mechanistic model of HCV entry. **A.** HCV particles must acquire CD81 to enter; this can be achieved via two routes, 1. acquisition of CD81 via intrinsic binding, with no prior involvement of other receptors 2. prior engagement of SR-B1 primes E2 for subsequent interaction with CD81. **B.** A parameterised mathematical model of virus entry. Each box represents the receptor engagement state of cell-attached viruses (*M*), where the values in parentheses denote receptor number. For example, *M* (0, 0) represents viruses that have attached but have not yet engaged either receptor, *M* (1, 0) represents viruses that have acquired one molecule of CD81 and *M* (0, 1) are viruses with one molecule of SR-B1. To achieve productive infection (*I*), virus particles must move laterally, acquiring CD81, this can be achieved via route 1 or 2, as described in A. The steps in this process have been parameterised to include the availability of receptor (determined by experimental data) and the rate of engagement (estimated by modelling). The dimensions of this matrix represent the stoichiometry of receptor engagement (i.e. how many molecules of CD81 are required for entry), we investigated this through the modelling process. There are two additional parameters in the model, e integrates all downstream events in the virus life cycle leading to productive infection, whereas *d* is the rate of spontaneous virus inactivation, or ‘death’. For clarity, parameterised steps are also shown in A.

Our model concerns the receptor engagement phase of entry, following virus particle attachment; acquisition of CD81 by HCV particles proceeds via two routes. The first utilises the intrinsic binding capacity of E2 for CD81 (Figure 4); in the absence of SR-B1 this pathway will allow a minority of particles to achieve entry (as demonstrated in Figure 2, S2). The second route to CD81 acquisition is via SR-B1. In this case, HCV particles gain SR-B1 through the strong intrinsic capacity of E2 to bind SR-B1 (Figure 4); this interaction confers an additional ability for HCV to acquire CD81, possibly through specific priming of E2-CD81 interactions. Thus, whilst SR-B1 is unnecessary for viral entry, it significantly increases the rate at which viruses enter the cell. Having acquired sufficient numbers of CD81, HCV particles progress to further stages of virus entry, including the acquisition of downstream receptors, endocytosis and fusion [4,34].

This putative entry mechanism was formulated as a mathematical model, which is summarised in Figure 5B. Viruses attached to the membrane are modelled as being in one of a number of states, *M* (*a, b*), representing a virus particle that has acquired *a* CD81 molecules and *b* SR-B1 molecules; upon initial attachment to the membrane the virus occupies the state *M* (0, 0). Subsequently, viruses are able to acquire molecules SR-B1 and CD81; initially, the rate of receptor engagement is determined by their intrinsic propensity for binding to these molecules, denoted by the parameters *s* and *c*_1_, respectively. As viruses bind SR-B1 a further pathway to CD81-acquisition becomes available, whereby viruses gain CD81 at rate *c*_2_. To integrate experimental data on receptor availability (Figures 2 and 3), the model allows CD81 and SR-B1 availability to be scaled by the terms *p*_*c*_ and *p*_*s*_, respectively, which are expressed relative to unmodified cells. For example, a value of 0.1 would be equivalent to a 90% reduction in receptor availability (via antibody blockade, Figure 2), whereas a value of 2 is achieved by doubling the amount of receptor (via over-expression, Figure 3). Having acquired a sufficient number of CD81 molecules to achieve entry, viruses proceed to cause infection, represented by entering the state *I*. This post-CD81 step occurs at the rate *e*: this parameter incorporates all downstream events leading to productive infection, including both the later stages of entry and the subsequent steps in the virus life cycle necessary for protein expression. Finally, to reflect the intrinsic instability of HCV particles [40] we also include a ‘death’ rate (*d*), this determines the rate at which viruses spontaneously inactivate and are therefore unable to achieve entry. Without loss of generality, the value of *d* was set to an arbitrary value, all other parameters being calculated relative to this. We constrained the model by inputting experimentally determined values for receptor availability and productive infection (Figures 2 and 3), and then optimised the parameters *s*, *e*, *c*_1_, and *c*_2_ so as to achieve a best fit model to the data obtained from our experiments. Data encompassing the effects of SR-B1 and CD81 receptor availability were fitted simultaneously.

Further to the above approach we also explored the effect of constraining the intrinsic binding rates to CD81 and SR-B1, described by the terms *c*_1_ and *s*. As discussed above, using the soluble E2 experiments to determine the ratio between *c*_1_ and *s* is inappropriate due to the limitations of the assay. However, given the observations in Figure 4, it is reasonable to expect that the rate at which HCV acquires SR-B1 (*s*) is somewhat greater than the rate of CD81 acquisition (*c*_1_). We denote the ratio between these values by *λ* = *c*_1_*/s*, for instance *λ* = 2 corresponds to SR-B1 acquisition occurring twice as fast as CD81 acquisition. In the case of an unconstrained fit to data we obtained a value of *λ* = 0.7. Increasing values of *λ* produced worse fits to the data, suggesting that this value is unlikely to be high (Supporting Figure S7). Therefore, given our combined observations, we estimate that a value of between 1 and 3 is most consistent with the observed data.

### Model outputs and predictions

Our model estimates the probability that a single virus founds a productive infection under varying availability of CD81 and SR-B1. Conversion of this statistic to the probability that a single cell is infected in the assay of Supporting Figure S1 can be obtained by accounting for the number of attached virus particles in the experiment (Figure 1).

Optimisation of model parameters gave an excellent fit to the experimental data (Figure 6). Inferences from our model further characterise the fate of virus particles: only 20-30% of HCV particles which attach to the membrane acquire a sufficient number of CD81 molecules to proceed with virus entry. Downstream events leading to productive infection are then even less efficient, with only 4-8% of particles completing these steps. Consequently, a virus particle participating in an infection assay has a very small chance of success: only *∼* 5% of particles attach to the membrane (Figure 1) with only *∼* 1.4% of these achieving productive infection (Table 1).

**Table 1.** **Optimised model outputs.** is table displays optimised values for the critical model parameters outlined in Figure 5. Rates are calculated relative to a viral death rate of 0.01. Values are shown for models optimised with *λ* = 1, 2, 3 and with no constraint placed on *λ*. In addition, the most likely CD81 stoichiometry derived from each model is given. The model can also be used to predict the fate of membrane bound virus particles. Here, we provide the inferred proportion of viruses that acquire sufficient CD81, that complete downstream steps and that achieve productive infection. Figures 6, 7 and 8 are generated using a model optimised with *λ* = 2. As such, the values from this model have been highlighted in bold.

| Optimised parameters | Unconstrained | $\lambda = 1$ | $\lambda = 2$ | $\lambda = 3$ |
| --- | --- | --- | --- | --- |
| Intrinsic ability to acquire SR-B1 ($s$) | 0.112 | 0.078 | **0.036** | 0.032 |
| Intrinsic ability to acquire CD81 ($c_1$) | 0.172 | 0.078 | **0.018** | 0.011 |
| Primed acquisition of CD81 ($c_2$) | 0.135 | 0.100 | **0.049** | 0.036 |
| Likely CD81 stoichiometry | >11 | 8 | **4** | 3 |
| Model predictions |  |  |  |  |
| Proportion of particles that acquire sufficient CD81 | 19.25% | 26.89% | **31.00%** | 32.71 % |
| Proportion of particles that complete downstream events | 7.30% | 5.29% | **4.57%** | 4.35 % |
| Proportion of particles that achieve infection | 1.40% | 1.42% | **1.42%** | 1.42 % |

**Figure 6.**
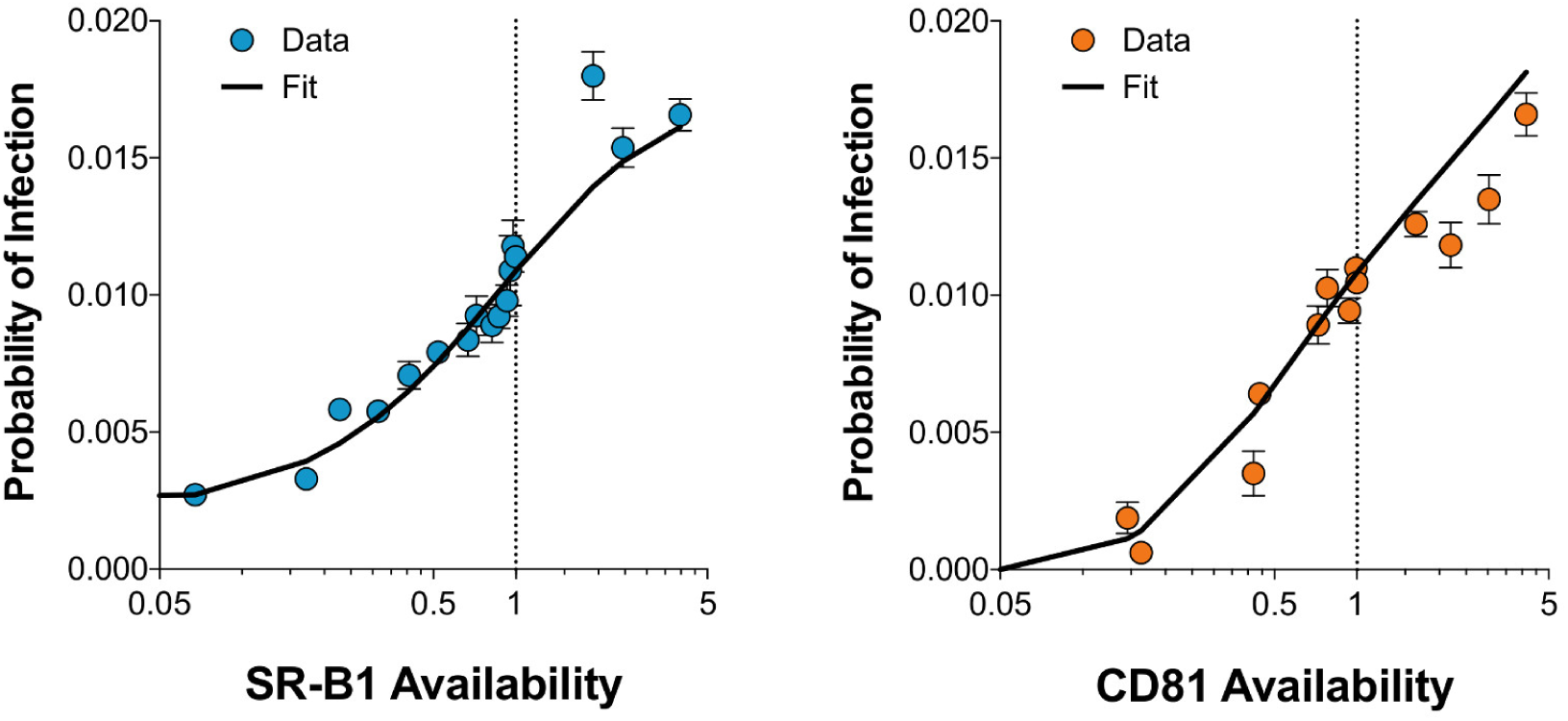
The mathematical model achieves excellent fit with experimental data. Multiple independent receptor blockade/over-expression experiments allowed us to determine the relationship between receptor availability and HCV infectivity. By integrating the virus attachment data from Figure 1 we estimated the probability of an attached virus yielding a productive infection for each condition. The X-axis represents receptor availability where values below 1 are derived from receptor blockade experiments and values above 1 are from over-expression experiments. The left Y-axis displays the probability of a cell within the experimental setup being infected by virus. The data points display mean probability of infection under varying receptor availability from *n* = 3 independent experiments (for both receptor blockade and over-expression, i.e. *n* = 6 experiments in total). Error bars indicate standard error of the mean. The black line displays the fit achieved with a model optimised with *λ* = 2; the data and fit are in excellent agreement.

Model predictions on the fate of virus particles were consistent across the range of tested *λ* constraints (Table 1). However, not all inferred parameters were robust to variations in this ratio. For example, the number of CD81 receptors required for entry was sensitive to the constraint placed on *λ* (Supporting Figure S8); as such our estimate of receptor stoichiometry is not as precise. With unconstrained *λ* no maximum number of required CD81 receptors could be found; increasingly small gains in likelihood were obtained with each additional receptor required by the model. In contrast to this result, *λ* values greater than 1 suggested lower receptor stoichiometry. Figure 7 displays the maximum likelihood (i.e. quality of model fit) for increasing numbers of CD81 molecules when *λ* = 2; a distinct peak in likelihood is observed at around 4 molecules of CD81. Given this, we suggest that 3-6 molecules of CD81 may be necessary and sufficient HCV entry.

**Figure 7.**
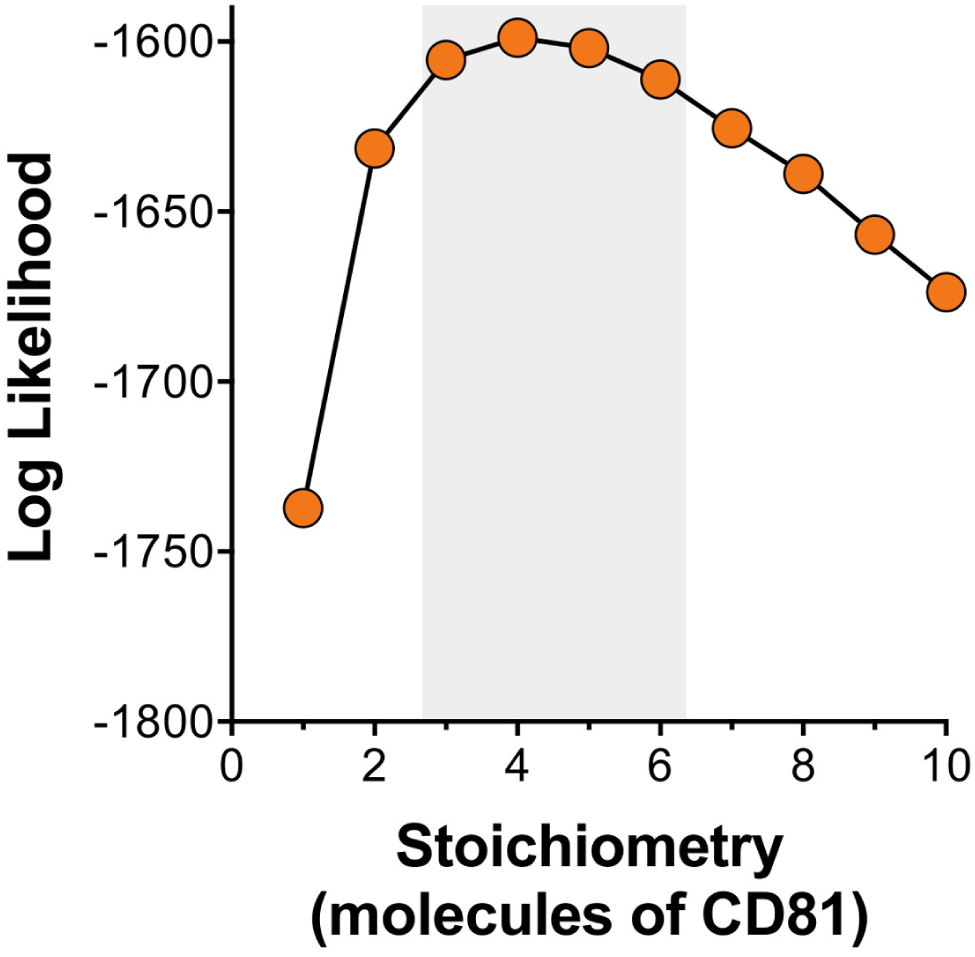
Predicting the stoichiometry of HCV-CD81 interactions. To investigate the number of CD81 molecules that are necessary for HCV entry we performed model optimisation across a range of potential HCV-CD81 stoichiometries. The log likelihood provides a measure of model fit, with higher values indicating a better fit. The data presented here was generated with a model optimised with *λ* = 2. A distinct peak in likelihood, indicating the best model fits, was observed at 3-6 molecules, as highlighted. This suggests a relatively low HCV-CD81 stoichiometry during virus entry.

### Investigating receptor interdependency

Our model predicts a specific interdependence in HCV receptor usage that we can confirm experimentally. We used the model to investigate the effect of co-varying SR-B1 and CD81 availability; in effect, this enables in silico infection assays to be performed with arbitrary receptor availability (Figure 8A).

**Figure 8.**
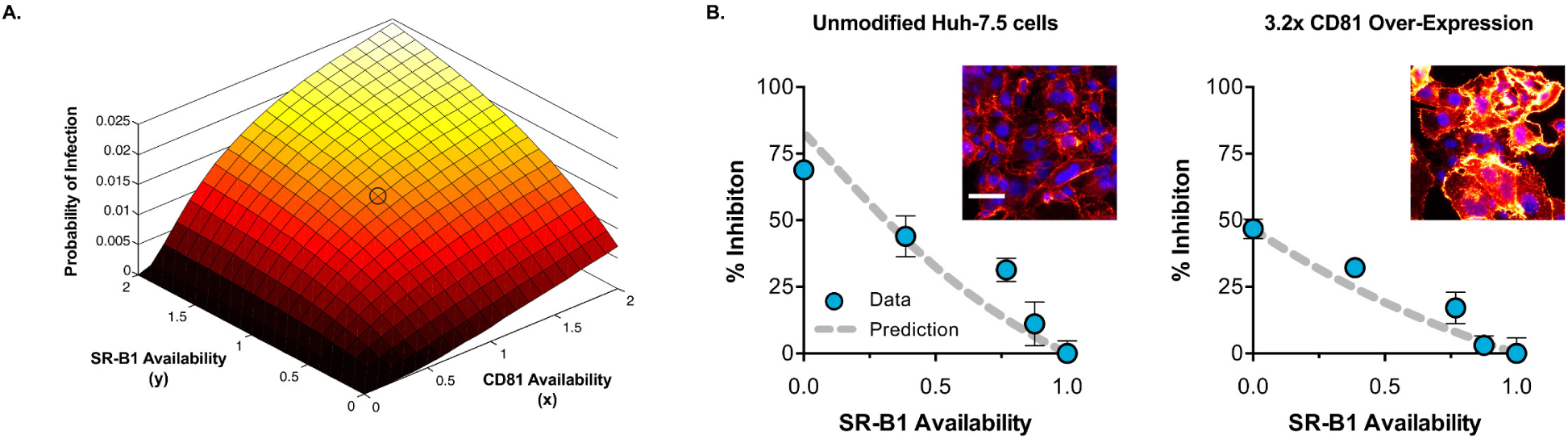
Experimental verification of model predictions. Our mechanistic model can be used to predict the probability of infection for any given availability of SR-B1 or CD81. **A.** Surface plot demonstrating the effect of co-varying receptor availability from 0-2, the circled area indicates the probability of infection in unmodified cells. Note that in the absence of CD81, altering SR-B1 levels has no effect; in contrast increased CD81 availability is able to rescue infectivity in cells lacking CD81. We experimentally tested this relationship between CD81 expression and SR-B1 dependence. **B.** Inset micrographs display CD81 and DAPI stained samples of unmodified Huh-7.5 cells or those transduced with lentivirus encoding CD81; 3.2x over-expression was achieved. Scale bar 50*μ*m. We investigated SR-B1 dependence in the over-expressing cells by antibody-mediated receptor blockade (as in Figure 2). To aid comparison, the data is expressed as % inhibition upon decreasing SR-B1 availability (compared to unblockaded cells). We used the model to predict the outcome of the experiment; in 3.2x CD81 over-expressing cells we expect that complete blockade of SR-B1 inhibits infection by *∼* 50%, compared to *∼* 80% inhibition for unmodified cells. Our experimental data is in excellent agreement with the model predictions. Data points represent mean values of *n* = 4 technical replicates, error bars indicate standard error of the mean. Data from one experiment are shown.

We evaluated the predictive power of this model via an experimental test of the effect of co-varying receptors. CD81 availability was increased by lentiviral transduction to achieve a ~3-fold increase in surface expression. These cells were then subjected to anti-SR-B1 receptor blockade, so as to evaluate SR-B1 dependence. Figure 8B displays the predicted outcome along with empirical measurements from this experiment. To aid comparison the data is presented as % inhibition upon decreasing SR-B1 availability, i.e. at normal SR-B1 availability infection is uninhibited and is therefore presented as 0% inhibition. As indicated in Figure 8A, the model predicts that high CD81 availability reduces the inhibitory effect of SR-B1 blockade; specifically, at *∼*3-fold CD81 over-expression *∼*50% of entry events occur in a SR-B1 independent manner, compared to *∼*20% in unmodified cells. Our empirical measurements are in excellent agreement with the predicted outcome of the experiment.

To summarise, we have used combined experimental and mathematical approaches to provide a number of mechanistic insights into the early processes of HCV entry. Both CD81 and SR-B1 are necessary for efficient entry, however, SR-B1 is somewhat dispensable and a minority of viral particles achieve entry in its absence (Figure 2 and S2). Soluble E2 binding assays suggest that HCV interactions with SR-B1 are more robust than those with CD81 (Figure 4). We provide a mechanistic model that can be used to understand these observations (Figure 5) and, through mathematical formulation and fitting, demonstrate that is model is consistent with our experimental data (Figure 6). This model was then used to make further predictions about the process of entry. Our work supports a relatively low stoichiometry of HCV-CD81 interactions during entry with less than or equal to six receptor molecules required for infection (Figure 7). Moreover, we predict specific interdependencies between SR-B1 and CD81 that can be confirmed experimentally (Figure 8), this provides further support to the veracity of our model.

## Discussion

We have here outlined a novel quantitative model of the early steps of HCV viral entry, based upon newly collected experimental data. By contrast to a previous model of viral entry [24] our approach accounts not only for the role of CD81 receptors in viral entry, but also the influence of SR-B1 receptors upon this process. Our accounting for this interaction is minimally complex, proposing SR-B1-mediated and non-SR-B1-mediated pathways for the acquisition of CD81 receptors, each occurring at different rates. Nevertheless our model produces an extremely good fit to the data collected. Validation of the model against independently generated experimental data shows its ability to predict in vitro rates of viral entry, providing increased confidence that our model is broadly correct.

Our model produces consistent estimates of the proportion of HCV particles achieving infection, showing that a very small proportion of viruses achieved the necessary steps of association, binding CD81 receptors, and the following downstream entry steps required to achieve infection. In our experiment only *∼*25% of particles that associated with cells completed the acquisition of CD81 receptors; having achieved this, only *∼* 5% of viruses then proceeded to productive infection. We expect that similar inefficiencies occur in vivo, and are likely to be compounded by the presence of non-permissive target tissues and an active immune response. Together the difficulties in viral entry may contribute to the extreme bottleneck observed for HCV transmission, where infection is established by only 1-5 viral particles [41,42].

While being less precise in this statistic, our model provides estimates of receptor stoichiometry, suggesting that between 3 and 6 CD81 molecules are required for infection. A previous mathematical modelling approach inferred that 1-13 E2-CD81 interactions being necessary for HCV entry [24]; our estimate is in broad agreement with this. Our model improves upon previous approaches in its use of a specific mechanistic model incorporating the gradual acquisition of receptors, considering the influence of SR-B1 upon viral entry and the potential for interdependencies between viral receptors. We note that the number of glycoprotein complexes per HCV particle remains unknown; our estimates of receptor stoichiometry provide a feasible lower bound for the number of functional E2 proteins per virus. This may suggest that unlike related flaviviruses such as Dengue, which possess 180 E proteins per particle, HCV glycoprotein incorporation is low, similar to HIV. This is supported by immunogold EM imaging of isolated HCV particles, which demonstrated limited detection of E2 [43].

The most significant limitation in our approach, relevant to the estimation of stoichiometry, is the difficulty in quantifying the relative abilities of HCV to acquire SR-B1 or CD81 receptors (*c* _1_ and *s* in the model). Our experiments evaluating the binding of sE2 binding suggest that E2 binding to SR-B1 is much more robust than to CD81. This finding is supported by previous reports [44] and is consistent with the current structural understanding of E2, which suggests that the SR-B1 binding site (HVR-1) is highly exposed on the surface of E2, whereas the CD81 binding site is likely to be shielded within the tertiary fold of the protein [45-47]. However, numerous technical caveats mean that it is inappropriate to extract a verbatim interpretation of these experiments. Conservatively, therefore, we conclude that the E2-SR-B1 interactions are somewhat greater than E2-CD81 (i.e. *c*_1_*/s >* 1), using the model to investigate this parameter further. Model fits to the experimental data then show a declining likelihood for values of this ratio that are much greater than 1 (Figure S7). We therefore explored a region of model space in which the ratio is not exceptionally high. The majority of the outputs presented here (Figures 6, 7 and 8) are generated from a model in which the ratio was constrained to 2, i.e. SR-B1 binding is twice that of CD81 binding. Accurate measurements of HCV particle-receptor interactions would likely improve the predictive power of our model.

Further limitations in our model can also be considered. Our model assumes a homogeneity of virus particles and cells, for example ignoring variation in receptor density between cells. This assumption may not be too problematic: our experimental data are ensemble measurements derived from a large number of viruses infecting a large number cells, as such, any heterogeneity is likely to be averaged out across these large populations. We further assume that each of the E2 glycoproteins on a virus particle acts independently, that is to say that receptor binding of one molecule of E2 does not positively or negatively impact the functionality of its neighbours. While this assumption may be problematic, speculation about alternative possibilities is difficult due to the limited information about the arrangement or oligomeric state of glycoproteins on HCV particles.

Our choice of model design, in which HCV achieves viral entry by the stepwise accumulation of receptors, is supported by a broad range of experimental evidence. For example, Baktash et. al. perform single particle tracking of HCV entry in to human hepatoma organoids [34]. This work demonstrates that entry occurs via a two-phase process, in the first of which HCV forms a tripartite receptor complex with SR-B1, CD81 and EGFR at the basolateral surface. Numerous other studies have assessed HCV entry kinetics using time of addition inhibitor studies; these experiments suggest that SR-B1 functions immediately after virus attachment, and likely precedes CD81 engagement [12,36,37]. Furthermore, multiple reports have been made of a minority fraction of infection that persists despite targeting of SR-B1 with CRISPR, siRNA or antibody blockade [35,37,48,49], supporting the notion of an SR-B1-independent mode of entry.

Our model provides new insight into the HCV entry process, suggesting that initial E2-SR-B1 interactions prime subsequent CD81 interactions. Mechanistically, this is most easily explained through analogy to HIV entry, where primary interactions between Env and CD4 stabilise an alternative conformation of gp120 that is activated for secondary binding to CCR5/CXCR4 [50,51]. This type of mechanism also fits with the presentation of the receptor binding sites on E2: SR-B1 interaction may directly or allosterically result in the unshielding of CD81 binding residues. However, despite ongoing investigations we have yet to gain direct evidence of SR-B1-dependent conformational changes in E2. Alternatively, SR-B1-dependent priming may occur in a less direct manner; for instance, the orientation of a virus particle bound to SR-B1 may promote CD81 binding. Indeed, recent crystallographic studies suggest that CD81 adopts a compact conformation that is likely to project from the plasma membrane by only 1-2nm [52]. In contrast the large ectodomain of SR-B1 is thought to form a barrel like conformation that extends much further from the cell surface [53].

This work provides a template for understanding the molecular processes of HCV entry and raises a number of pertinent questions for future investigation. For instance, the receptor binding capacity of E2 is likely to vary under viral adaptation, therefore, we may expect different HCV strains to exhibit different entry characteristics. Our modelling approach could be used as an investigative tool to ask important questions in this area, for example: do the entry characteristics of transmitted/founder viruses differ from those circulating in a chronically infected person. Furthermore, our work provides evidence for sequential, interdependent receptor interactions; are these functional stages of virus entry determined by specific conformations of the viral glycoproteins, and if so, which of these conformations is most relevant for B-cell immunogen development.

## Methods

### Experimental methods

#### Cell lines

Huh-7.5 cells were provided by APATH LLC; Huh-7 and CRISPR Cas9 receptor KO cells were a gift from Yoshiharu Matsuura (Osaka University) [35]; CHO and HEK 293T cells were acquired from the American Type Culture Collection. All cells were propagated in Dulbecco?s Modified Eagle Medium (DMEM) + 10% fetal calf serum (FCS) supplemented with penicillin, streptomycin and non-essential amino acids (Life Technologies, Carlsbad, CA, USA).

#### Antibodies

Rabbit anti-SR-B1 polyclonal serum was a gift from Thierry Huby (INSERM, Paris); mouse monoclonals anti-CD81 (2.131) and anti-NS5 (S38) were gifts from Jane McKeating (University of Oxford); StrepMAB-Classic was acquired from IBA lifesciences (Gttingen, Germany).

#### Generation of HCVcc and infection assay

Full-length HCVcc RNA genomes were generated by in vitro transcription from J6/JFH plasmid template (provided by APATH LLC) [26,33]. To initiate infection, viral RNA was electroporated into Huh-7.5 cells using a BTX830 (Harvard Instruments, Cambridge, UK). From 3-7 days post electroporation, cell culture supernatants containing infectious J6/JFH HCVcc were harvested every 2-4 h; these short harvest times limit the opportunity for virus degradation, therefore, preventing the accumulation of non-infectious particles. A uniform experimental stock of virus was generated by pooling the harvested supernatants; this ensures maximum reproducibility between experiments.

To assess HCVcc infectivity Huh-7.5 cells were seeded at a density of 1.5 *×*10;^4^ in to each well of a standard flat bottomed 96-well tissue culture test plate 24 h prior to study. Receptor availability was modulated as described below. To infect, the cells were refed with 50*μ*l of DMEM + 3% FCS and challenged with an equal volume of HCVcc supernatants diluted 1/2 in DMEM + 3% FCS (this resulted in each well receiving *∼*180 foci forming units of infectious virus). After 5 h the inoculum was removed, and the cells were washed twice in 50 *μ*l PBS and refed with 100*μ*l DMEM 3% FCS. Cells were fixed after 48 hours by washing with 50*μ*l of PBS and treatment with 250*μ*l ice cold methanol for 10 min. To detect viral antigen, samples were blocked with 0.5% bovine serum albumin (BSA) stained with anti-NS5 (S38) hybridoma supernatant (1/100), followed by 2 *μ*g/ml goat anti-mouse Alexa Fluor 647 secondary antibody (Life Technologies). Nuclear DNA was counterstained with 2 *μ*g/ml 4’,6-diamidino-2-phenylindole (DAPI).

To quantify infection assay plates were imaged using a Nikon Ti inverted microscope fitted with a motorized encoded stage for plate-reading. A 4 mm by 4 mm area of each well was acquired by image stitching using an ORCA Flash 4 sCMOS camera (Hamamatsu, Welwyn Garden City, UK), with 405 nm and 647 nm fluorescence illumination provided by a PE4000 LED unit (CoolLED, Andover, UK) through a multi-band excitation/emission filter cube (Semrock, Rochester, NY, US). To ensure optimal imaging, software-based autofocusing was performed prior to acquiring each well. The number of foci in each image was quantified manually using the counting tool in FIJI/ImageJ [54,55]. In addition, the number of DAPI nuclei were counted using the ‘find maxima’ function, this data was later used to derive estimates of the probability of infection (described below).

#### Antibody-mediated receptor blockade

To limit the availability of receptor, Huh-7.5 cells seeded for infection were pre-incubated at 37C with 50*μ*l of media containing a serial dilution of either rabbit anti-SR-B1 polyclonal serum [33] or mouse anti-CD81 (2.131) hybridoma supernatant [32]. After 45 mins, the wells were challenged with virus, as described above.

#### Lentivirus-mediated over-expression

To generate lentiviral vectors we transfected HEK 293T cells with three plasmids: a HIV packaging construct (pCMV-dR8.91), VSV-G envelope plasmid (pMD2.G) and a dual promoter transfer plasmid encoding GFP in combination with SR-B1 or CD81 (pDual SR-B1 or CD81, available from Addgene). Huh-7.5 cells were transduced with lentivirus vectors, diluted in DMEM + 10% FCS, 96 hours before an experiment; 24 hours prior to study the cells were seeded into a 96 well plate for infection, as described above.

#### Quantification of receptor availability

In parallel with the infection assay we also prepared plates to assess receptor availability. Pre-treatment of these plates was identical to those being infected, however, instead of receiving virus they were mock inoculated with DMEM + 3% FCS. After 5 hours the plates were then washed once with PBS and fixed with 4% formaldehyde. The plates were then blocked for 1 hour in 0.5% BSA before staining to assess receptor modulation. To evaluate antibody-mediated receptor blockade, bound antibody was detected by addition of goat anti-mouse/rabbit Alexa Fluor 647 secondary antibody. To evaluate receptor over-expression, fixed cells were incubated with a saturating concentration of anti-SR-B1 or CD81 followed by anti-mouse/rabbit Alexa Fluor 647 secondary antibody. Nuclear DNA was counterstained with 2 *μ*g/mL DAPI. Imaging was performed as described above and image analysis/quantification was performed in FIJI/ImageJ.

#### Quantification of viral copy number by qPCR

Virus particle attachment was assessed by qPCR quantification of cell-associated RNA genome copy numbers. Cells were seeded and inoculated as described above for the infection assay, however, after 5 hours the cells were washed and then lysed for viral RNA extraction using a QIAamp Viral RNA Mini Kit (Qiagen, Hilden, Germany). RNA quantification was performed using a Luna Universal qRT-PCR kit (New England Biolabs, Ipswich, MA, USA) with primers targeting the 5’ UTR of the HCV genome (sense primer: GCGAAAGGCCTTGTGGTACT, anti-sense primer: CACGGTCTACGAGACCTCCC) as previously described [56]. RNA copy number was determined by including a standard curve of in vitro transcribed HCV RNA in each qPCR run. Reactions were run on a CFX96 touch thermal cycler (Bio-Rad, Hercules, CA, USA).

#### Soluble E2 binding assay

A truncated, soluble version of the E2 glycoprotein (sE2) was cloned by PCR amplification from a plasmid encoding full length HCV genome and ligated into a vector encoding an N-terminal tPA secretion signal and a C-terminal Twin-Strep-Tag [57,58]. sE2 was produced by transient transfection of HEK 293T cells followed by protein purification using the Step-TactinXT system (IBA lifesciences, Gttingen, Germany). To assess E2 interaction with receptors, CHO cells were transduced with lentivirus encoding human SR-B1 or CD81, transgene expression was allowed for at least 72 hours before the cells were used in a sE2 binding assay, as previously described [58]. Briefly, cells were trypsinised into a single cell suspension, and then incubated in blocking buffer (1% BSA + 0.01% NaN3) for 20 minutes, note that the inclusion of NaN3 stops cellular metabolism and arrests internalisation of surface bound ligands. sE2, diluted in blocking buffer, was incubated with the cells for 1 hour at 37C, followed by two PBS washes. Bound E2 was detected using anti-Strep-Tag antibody and goat anti-mouse Alexa Fluor 647 secondary antibody. Samples were quantified on a LSR-Fortessa cell analyser and the data anlysed using FlowJo (both Becton Dickinson, Franklin Lakes, NJ, USA).

### Computational methods

Multiple independent receptor blockade and over-expression studies were performed, as illustrated in Figures 2 and 3. These, along with the quantification of particle attachment (Figure 1), provide the necessary information to estimate the effect of receptor availability on the probability of infection; this data can then be used to fit our model. Achieving these estimates required some analysis and processing of the raw data.

#### Processing of fluorescence data

The use of fluorescence data to quantify the number of receptors available for binding poses two challenges. Firstly, measurements of fluorescence are imprecise, with an unknown level of variance in a repeat measurement. Secondly, measurements collected from different cell populations may not give consistent results; the level of fluorescence indicating saturation may vary from one population to another. Measurements were therefore processed to produce normalised estimates of how many receptors were available in each population.

Firstly, to account for imprecision in fluorescence measurements, we clustered data from populations which had been titrated with different quantities of antibody, identifying sets of readings which could be meaningfully distinguished from each other. For each set of cells i having a constant antibody concentration we denote the fluorescence values collected as {*f*_*i*_}. Sorting each set of values, we carried out a two-sample T-test on all pairs of differing sets {*f*_*i*_} and {*f*_*j*_}, identifying the pair of sets for which the test produced the highest p-value. If this p-value was greater than 0.05, indicating that the two sets of values could not be clearly statistically separated, the two sets were merged, repeating the calculation of T-tests across the new pairs of sets. Where sets contained different numbers of fluorescence values, 100 random samples were drawn from the larger set, each of size equal to that of the smaller set, and the mean p-value across these samples was calculated. Examples of mean and standard deviations of fluorescence values from distinct antibody concentrations taken before and after this clustering procedure are shown in Supporting Figure S9, this relates to the raw data shown in Figure 2.

Secondly, clustered fluorescence data were normalised to estimate the proportion of receptor available in each set following the addition of antibody. For each set, the maximum fluorescence, denoted max *f*, was calculated as the mean of the fluorescence values from the cells with the highest amount of antibody; this was assumed to indicate saturation of receptors. Similarly, the minimum fluorescence, denoted min *f*, was calculated as the mean of the fluorescence values from the cells for which no antibody was added. For the set {*f*_*i*_}, the proportion of receptors available was then calculated as

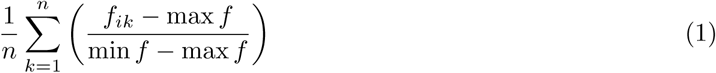

#### Processing of cell count data

Within the experiment, measurements of the total number of cells in each well were collected after 5 and 48 hours (Supporting Figure S1). These values were compared in order to account for the growth of cells during the experiment. This ratio was then used to scale the 48-hour cell counts, producing estimated 6-hour cell counts, corresponding to the number of cells in a well at the time of viral infection. We denote these estimated number of cells for a given population as *a*_*i*_.

#### Viral inoculum dose

An estimation of the effective viral dose for each set of cells was calculated. Attachment to the cell membrane is an essential precursor to viral entry. In measuring viral entry our concern is for events subsequent to this attachment. As such, rather than considering the pure ratio of number of viruses to number of cells in a well, we evaluated the mean number of viruses attaching to the cell membrane of each cell.

For a replicate set of cells, a large number of viruses was added to each well, following which the number of viruses bound to the membrane after 5 hours was calculated (Figure 1). This suggested that a mean of *n* = 11125 viruses were bound to the cell membrane. Cell counts were made using images which covered a 4mm_;^2^_; portion of each well, equivalent to a fraction of 0.454 of the well area. As such, the effective dose was calculated as

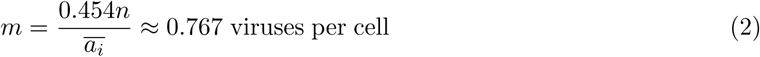

where the bar indicates a mean value across all populations. The dose was assumed to remain constant across all wells, assuming that small changes in the number of cells would be reflected by corresponding changes in the number of membrane-attached viruses.

#### Likelihood model

Given an a viral inoculum dose, the expected fraction *q*_*k*_ of cells in each well that are infected by *k* viruses, was modelled using a Poisson distribution:

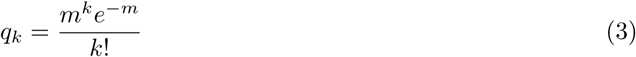

We developed a deterministic model, described below, to estimate the probability *P*_*i*_ that a virus which is bound to the cell membrane in well *i* gains viral entry. Supposing that viruses act independently of one another in gaining entry to a cell, the probability that at least one virus gains entry to a cell to which k viruses are bound is given by

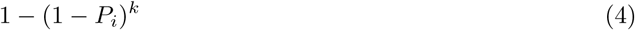

The number of foci in any given well is small relative to the number of cells. As such, we used a Poisson-like model to calculate the likelihood of our model. The total rate at which viruses gain entry into cells in well i is given by

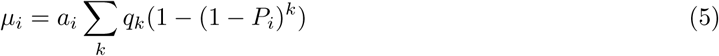

Given counts *o*_*i*_ of the number of foci of infection in each population, we then used a double Poisson model to calculate the likelihood of a model:

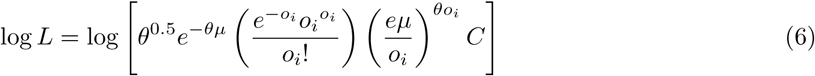

where

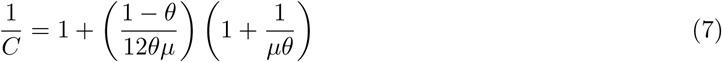

This model, while superficially similar to a Poisson model, grants additional flexibility, the variance of the distribution being controlled by the parameter *θ*. This parameter was estimated using a likelihood model to fit a probability *P*_*i*_ to data from each set of wells defined by the fluorescence data, without these probabilities being constrained by any underlying model. This gave the estimated value *θ ≈* 0.363, representing an increased variance in the data relative to a Poisson distribution.

#### Processing of overexpression data

Data were collected both from populations of cells grown with antibody to reduce the number of available receptors, and for populations of cells in which the over-expression of receptors was induced. To account for potential differences in cell populations, a linear scaling was applied to produce identical results for wild-type cells. Inferred frequencies from the estimation of were used to scale the parameters ai in the over-expression data to produce identical estimates of *P*_*i*_ for each equivalent set of wild-type cells; our model therefore assessed how changes in receptor availability lead to changes in viral entry.

#### Intra-cellular model of infection

In the framework above we require an estimate for the value *P*, describing the probability that a single virus that binds to the cell membrane gains entry to the cell. To calculate this value we use an ODE model of viral dynamics (Figure 5B) to estimate this. As described in the main text, our model contains the parameters *r*, describing the number of CD81 receptors required to gain entry into the cell, *e*, the downstream rate of viral entry having bound sufficient CD81 receptor, *s*, the rate at which viruses acquire SR-B1 receptors, *c*_1_ and *c*_2_, describing SR-B1-independent and SR-B1-mediated rates of acquiring CD81 receptors, and a ‘death rate’ for viruses, *d*.

Mathematically, the model can be expressed as follows:

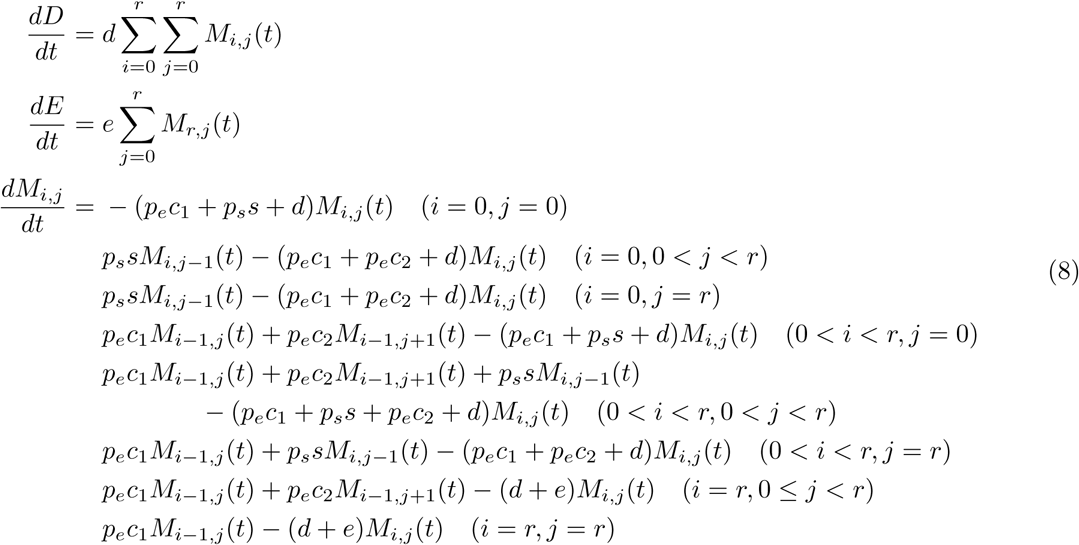

with initial conditions

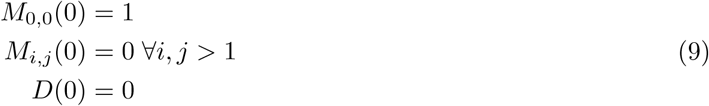

Simulations were run using a fourth-order Runge-Kutta scheme with adaptive step size until *D*(*t*) +*E*(*t*) *≥* 0.999. At this point, we calculate the value

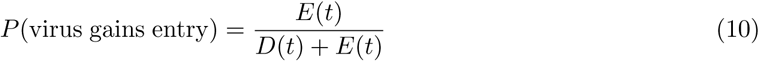

giving the proportion of viruses to have gained entry into the cell. Our model was run in an unconstrained form, and under a range of constrained values of the parameter *λ*, defined as the ratio *s/c*_1_. The death rate *d* was arbitrarily fixed to the value 0.01, all other rates being calculated relative to this. Likelihood values were compared for models in which different numbers of CD81 receptors were required for entry, specified by the parameter *r*.

#### Statistics of viral entry

Among the viruses which acquire sufficient CD81 receptors, not all go on to gain entry to the cell. The proportion of viruses which do gain entry can be simply inferred from the model parameters; having acquired sufficient receptors, virus particles gain viral entry at rate e, and die at rate d. The proportion of viruses with sufficient CD81 receptors which gain viral entry is therefore given by

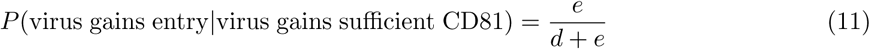

From this statistic we can derive the number of viruses which gain sufficient CD81 receptors to be able to potentially gain viral entry, as

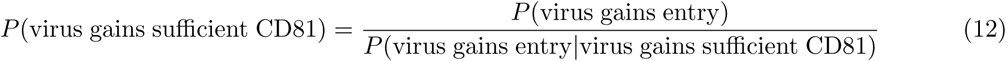

## Acknowledgements

This work was funded by The Wellcome Trust and The Royal Society; grant numbers 101239/Z/13/Z (CJRI) and 107653/Z/15/Z (JG).

## Availability of code

Code used in this research is available from https://github.com/cjri/HCVModel

## Supporting Figures

**Figure S1.**
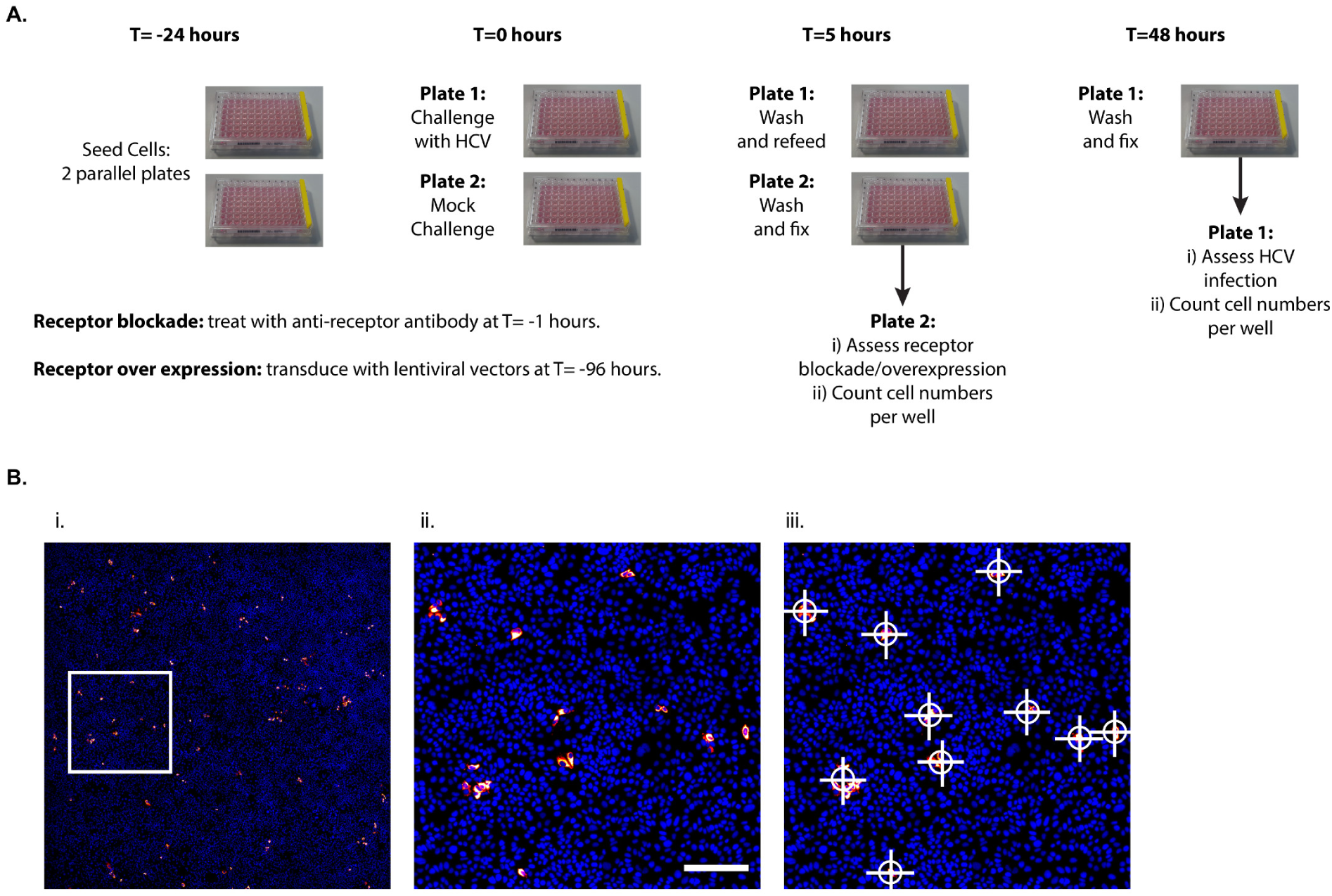
Infection assay workflow and quantification. **A.** The infection assay workflow: cells were manipulated to achieve receptor blockade or over-expression at the indicated time points. Plate 1 was used to determine HCV infectivity, whereas the parallel plate 2 was used to assess receptor blockade/over expression. **B.** Infection was quantified manually by counting infected foci at 48 hours. i. An example field of HCV infected Huh-7.5 cells stained for viral antigen and cellular nuclei. ii. A large image of the inset from A., multiple distinct foci of infection are apparent. iii. Viral infection was quantified by manual scoring of individual foci, as annotated on to the image. Scale bar 200*μ*m.

**Figure S2.**
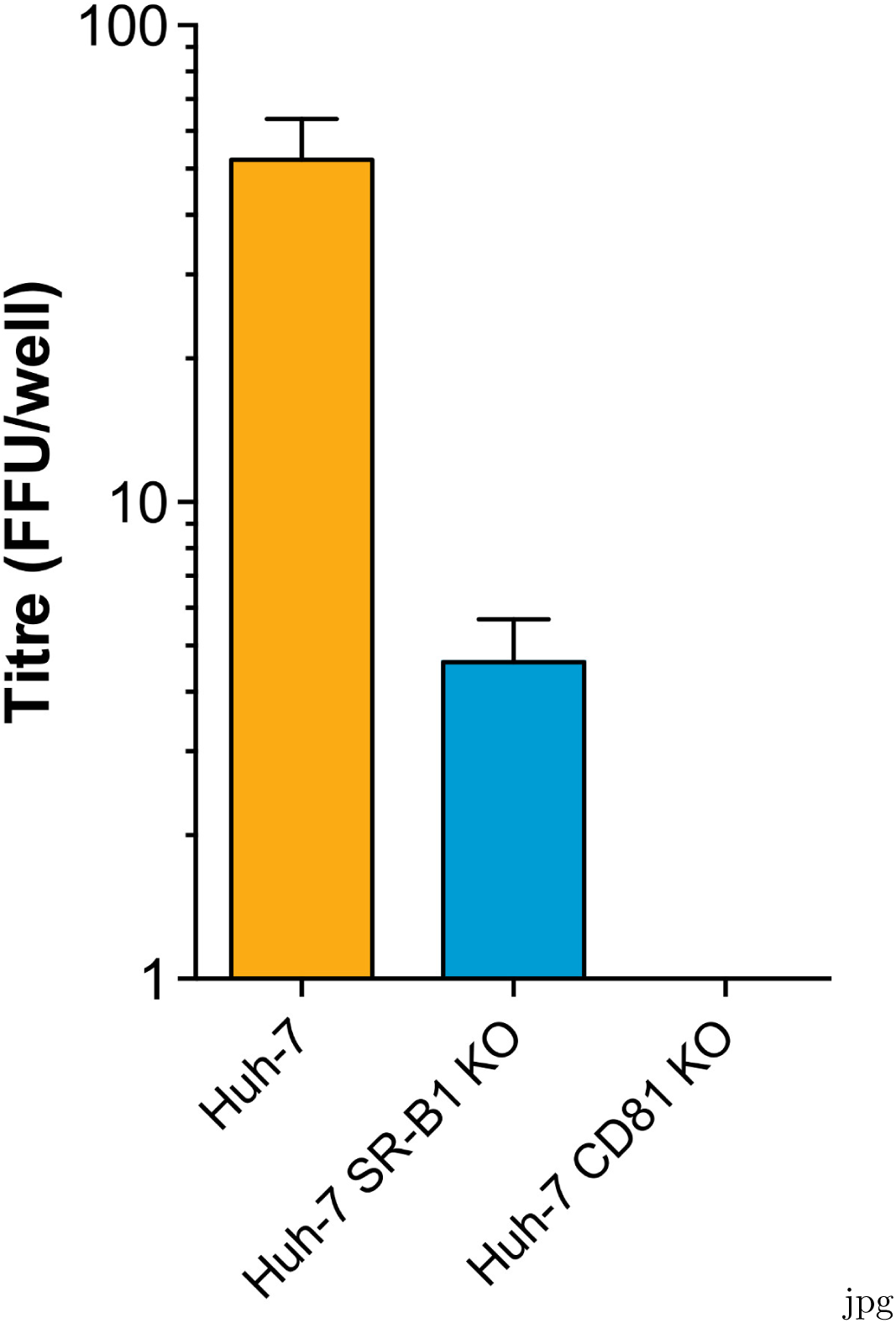
HCV challenge of receptor KO cells confirms SR-B1 independent infection. HCV titre in parental Huh-7 human hepatoma cells, or those in which receptor encoding genes have been knocked out by CRISPR Cas9 editing. Mean values of *n* = 3 independent experiments are shown. Error bars indicate standard error of the mean.

**Figure S3.**
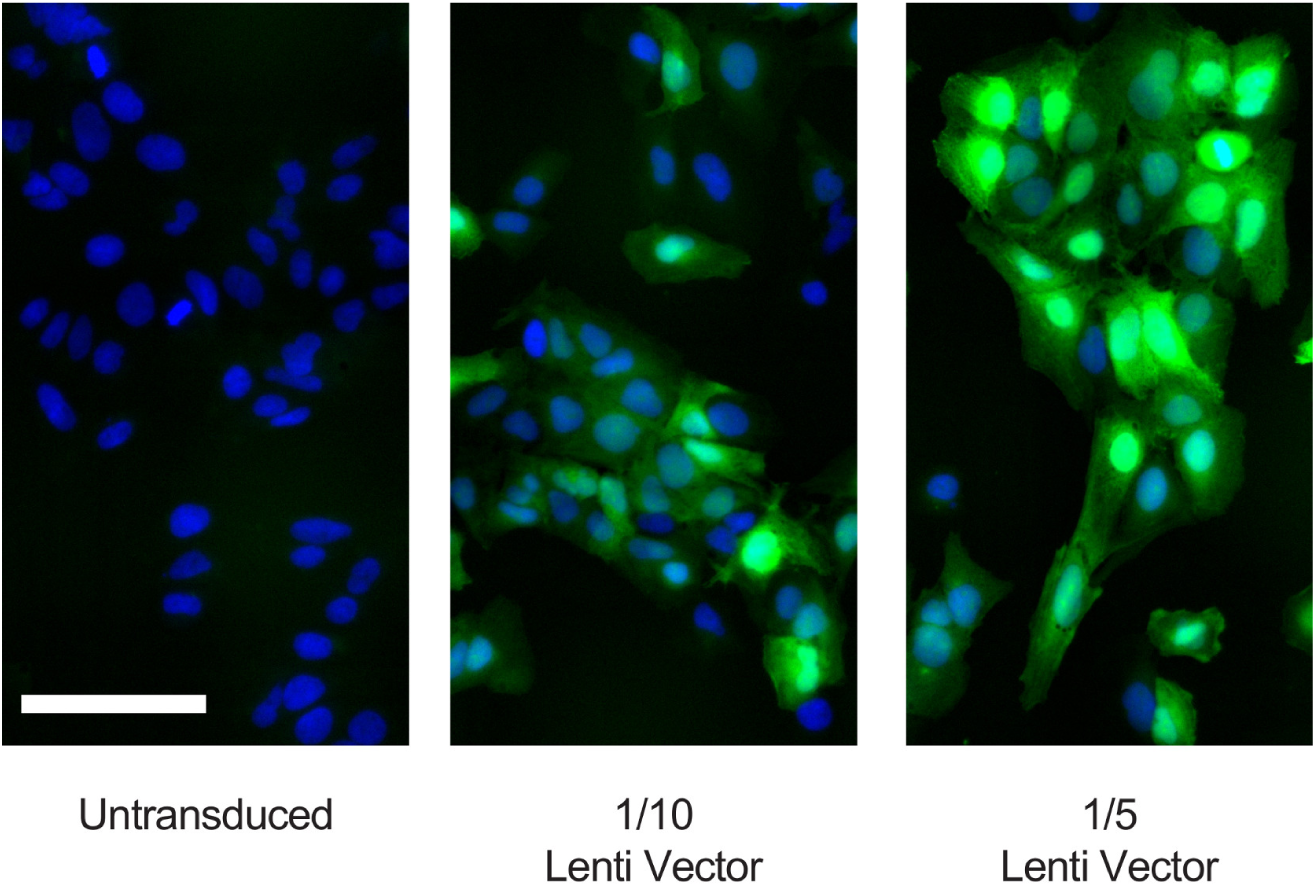
Lentiviral transduction of Huh-7.5 cells is homogenous. Huh-7.5 cells were transduced with lentiviral vectors that encode both a receptor (either SR-B1 or CD81) and GFP, expressed from separate promoters. Therefore, evaluating GFP expression provides an independent measure of transduction efficiency. The images display representative fluorescent micrographs of parental cells or those transduced with SR-B1 + GFP lentiviral vectors. GFP expression is homogenous between cells and titrates with lentivirus concentration.

**Figure S4.**
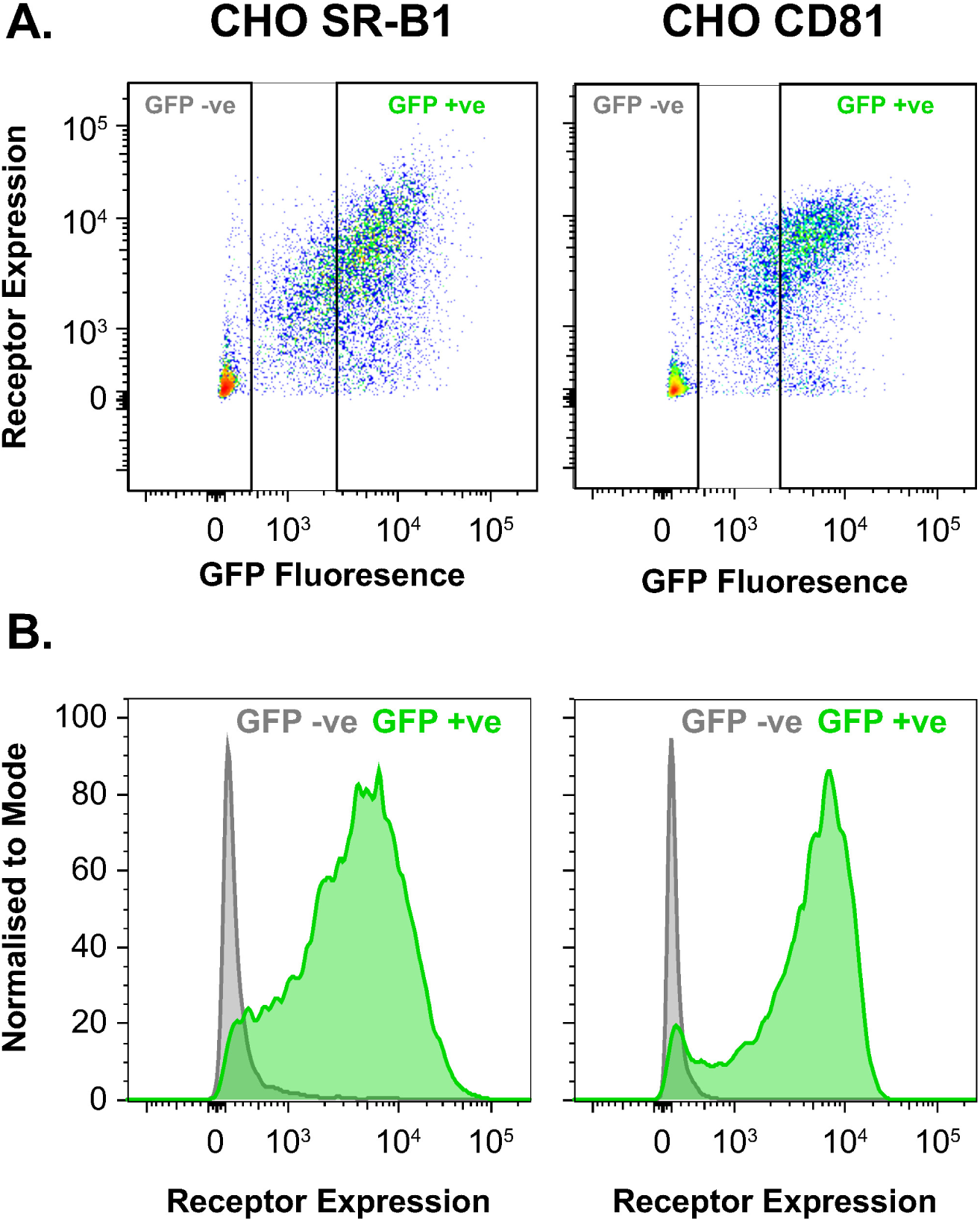
Transduced CHO cells express exogenous SR-B1/CD81. CHO cells were transduced with lentivirus encoding either SR-B1 or CD81 and GFP (as described in S3 Fig), receptor expression was assessed by flow cytometry. **A.** Representative dot plots of receptor and GFP expression in CHO cells, unlike Huh-7.5 cells, a minority of cells remained GFP/receptor negative. **B.** Representative histograms of receptor expression in GFP negative and positive CHO cells, as expected, receptor expression is only apparent in GFP positive cells.

**Figure S5.**
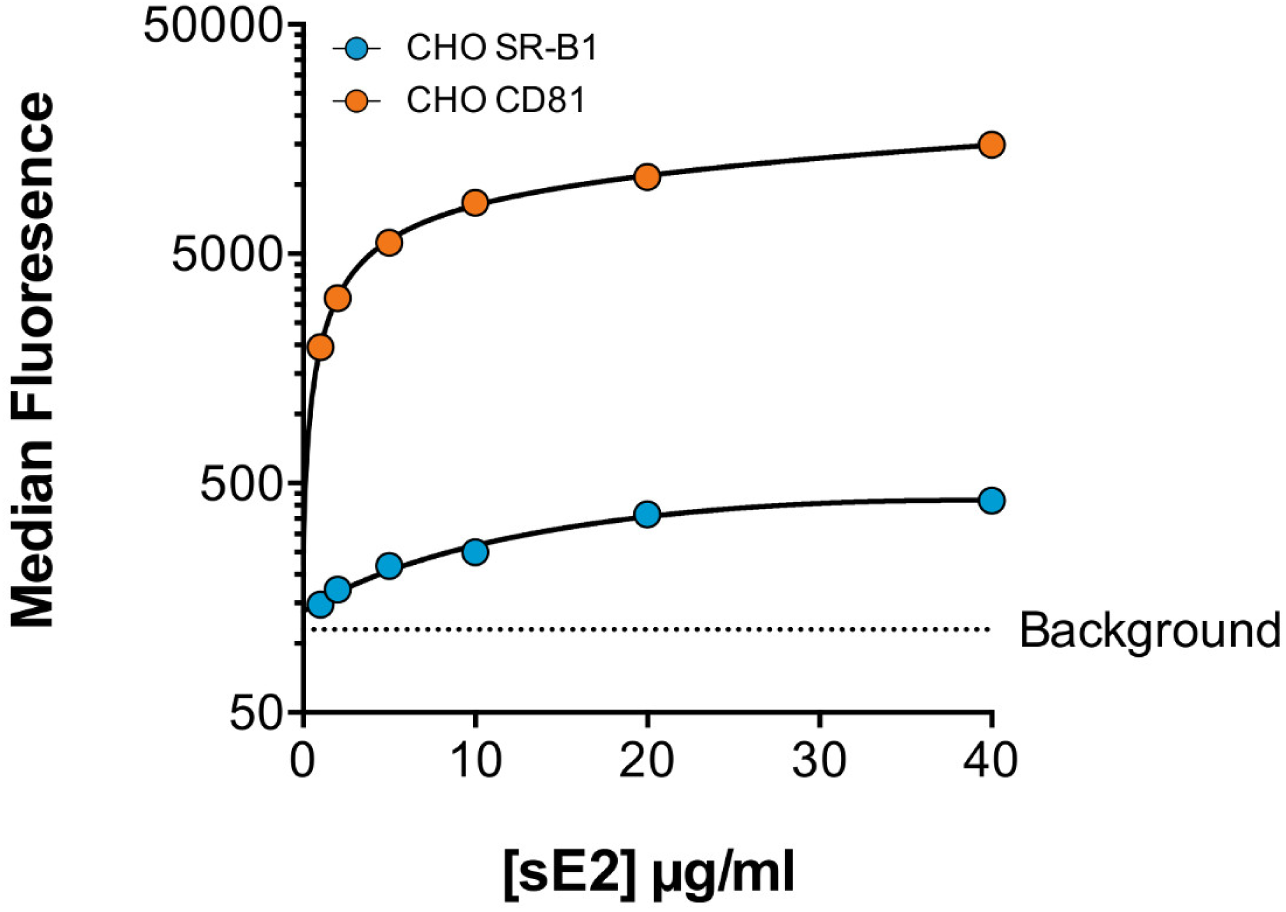
Representative raw data of sE2 binding to CHO SR-B1/CD81 cells. Representative median fluorescence intensity values for sE2 binding to CHO SR-B1/CD81 cells, as assessed by flow cytometry. Background is determined by sE2 binding to untransduced CHO cells. Data points represent the mean of *n* = 2 technical repeats. Error bars indicate standard error of the mean. Data was fitted using a one-site binding curve in GraphPad Prism.

**Figure S6.**
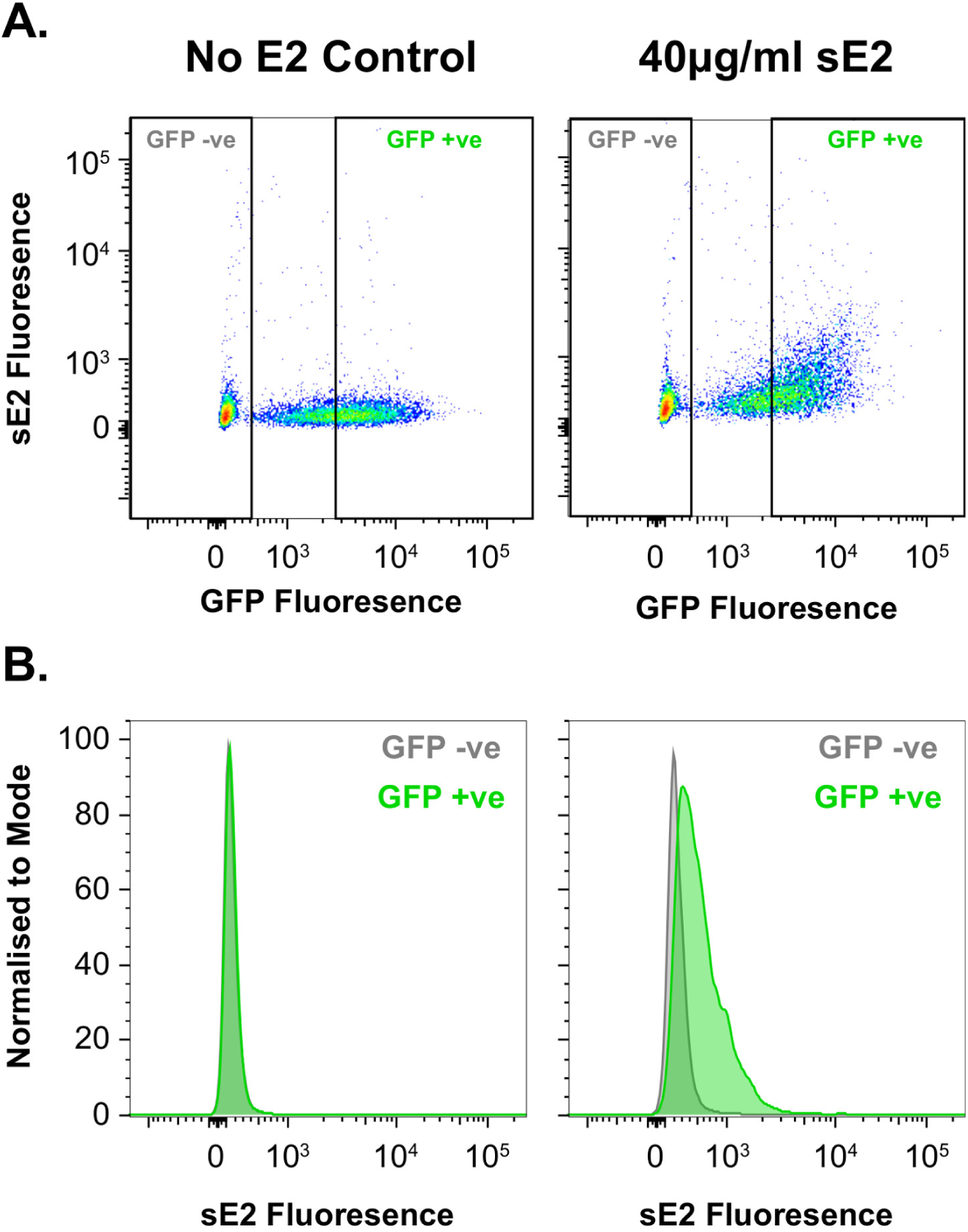
Soluble E2 binding to CHO cells expressing CD81 is low but readily detectable. Representative raw data showing sE2 binding to CHO cells transduced with lentiviral vectors encoding CD81 + GFP. **A.** Dot plots displaying sE2 binding and GFP expression in untreated CHO-CD81 cells and those incubated with 40*μ*g/ml sE2. **B.** sE2 binding to GFP negative and positive cells within the same sample, as expected, sE2 binding is only detectable in GFP positive cells, i.e those that have been successfully transduced with receptor encoding lentivirus.

**Figure S7.**
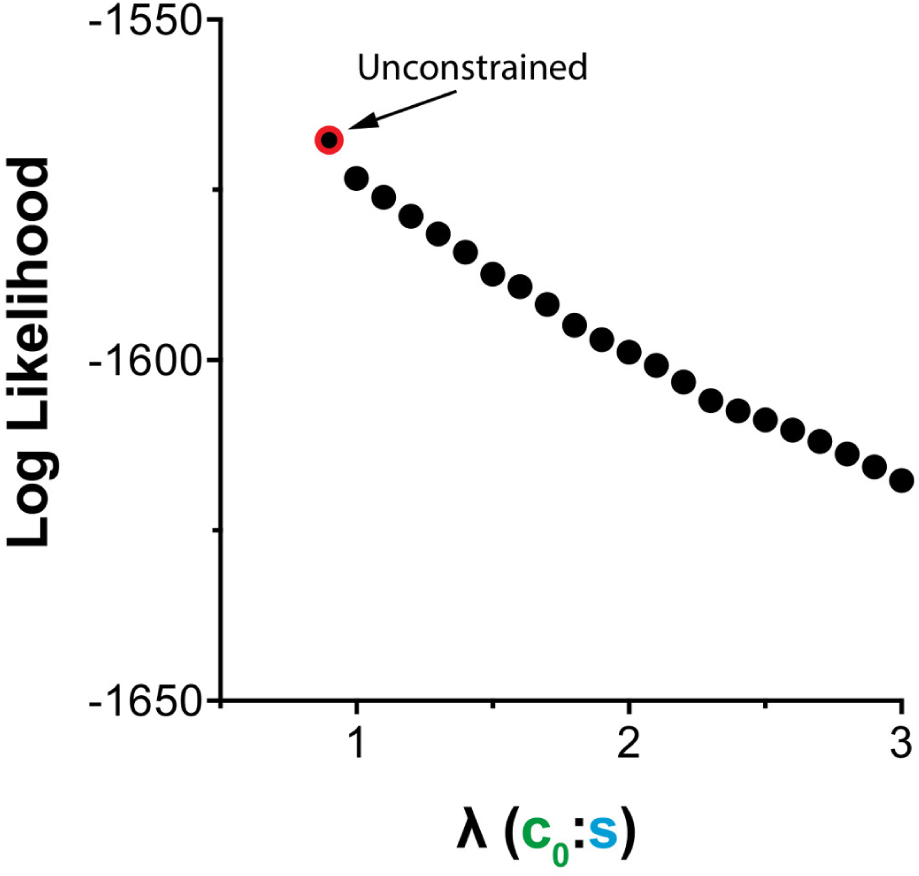
Increasing the *λ* constraint reduces the maximum likelihood of the model. sE2 binding data suggests that the intrinsic ability of HCV to acquire SR-B1 is greater than its ability to acquire CD81. We investigated this during model fitting by constraining the ratio between *c*_1_ and *s*, we denote this ratio as *λ*. This plot displays the maximum log likelihood for models optimised with different *λ* constraints; log likelihood provides a measure of how well a particular model fits the experimental data. Increasing values of *λ* resulted in a drop in log likelihood, this indicates that a high ratio is unlikely.

**Figure S8.**
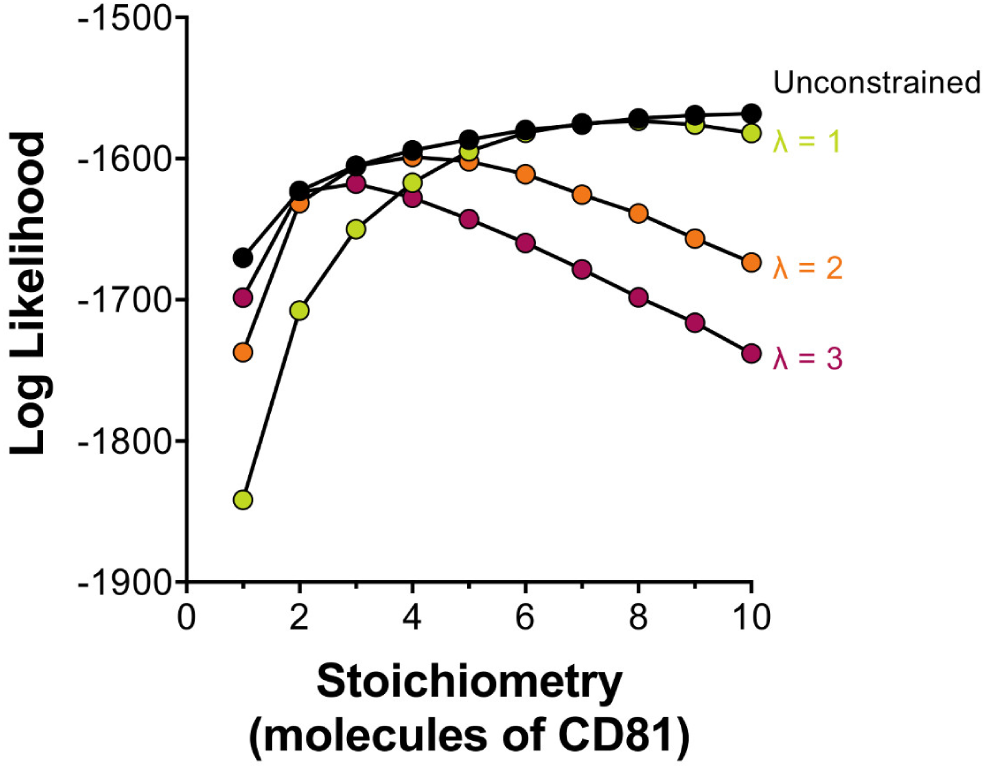
The inferred stoichiometry of HCV-CD81 interaction varies for models optimised with different *λ* values. To investigate the potential stoichiometry of HCV entry we varied the number of CD81 molecules necessary for entry during model optimisation. This plot displays log likelihood for models across a range molecule numbers; a peak in likelihood allows inference of the probable stoichiometry of HCV entry. When optimised without constraint placed on *λ*, we observed no peak in likelihood; each additional molecule resulted in an incrementally higher likelihood. However, models in which *λ* was constrained resulted in peak likelihoods at low stoichiometries of 2-6 molecules of CD81. This would suggest that relatively low numbers of CD81 are necessary for HCV entry.

**Figure S9.**
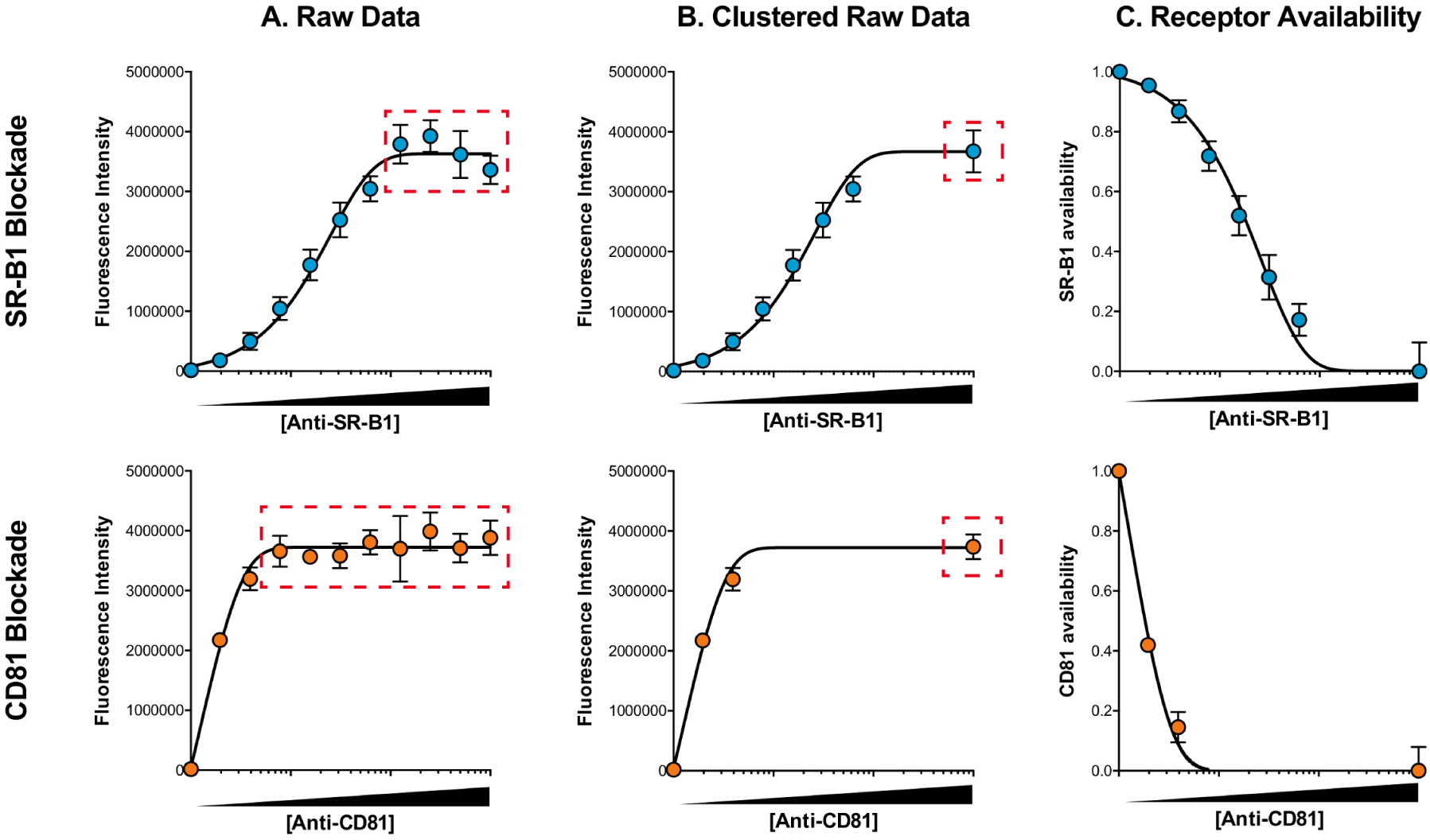
Clustering and processing of fluorescence data. We estimated receptor availability from fluorescence microscopy data. **A.** Representative raw fluorescence measurements of anti-SR-B1/CD81 binding to Huh-7.5 cells (similar to Figure 2A), data points represent the mean of n¿4 technical repeats, error bars indicate standard deviation of the mean. **B.** Statistically indistinguishable data points were clustered together and averaged; the clustered measurements and their resulting combined data point are annotated. **C.** Scaled estimates of receptor availability derived from clustered data. Maximum antibody binding indicates saturation of receptor and, therefore, availability equals 0; whereas in untreated cells receptor availability is set to 1. These receptor availability values can then be compared to matched infection data to explore HCV receptor availability. In each plot data was fitted using a sigmoidal curve in GraphPad Prism.

